# Coordinated development of the mouse extrahepatic bile duct: implications for neonatal susceptibility to biliary injury

**DOI:** 10.1101/576256

**Authors:** Gauri Khandekar, Jessica Llewellyn, Alyssa Kriegermeier, Orith Waisbourd-Zinman, Nicolette Johnson, Yu Du, Roquibat Giwa, Xiao Liu, Tatiana Kisseleva, Pierre A. Russo, Neil D. Theise, Rebecca G. Wells

**Author notes:** These authors contributed equally (Gauri Khandekar, Jessica Llewellyn). **ADDRESS CORRESPONDENCE AND REPRINT REQUESTS TO:** Rebecca G. Wells, MD, Department of Medicine (GI), Perelman School of Medicine, University of Pennsylvania, 905 BRB II/III, 421 Curie Blvd., Philadelphia, PA 19104-6140.

## Abstract

**Background & Aims:** The extrahepatic bile duct is the primary tissue initially affected by the cholangiopathy biliary atresia. Biliary atresia affects neonates exclusively and current animal models suggest that the developing bile duct is uniquely susceptible to damage. In this study, we aimed to define the anatomical and functional differences between the neonatal and adult mouse extrahepatic bile ducts.

**Methods:** We studied mouse passaged cholangiocytes, mouse BALB/c neonatal and adult primary cholangiocytes and isolated extrahepatic bile ducts, and a collagen reporter mouse. Methods included transmission electron microscopy, lectin staining, immunostaining, rhodamine uptake assays, bile acid toxicity assays, and in vitro modeling of the matrix.

**Results:** The cholangiocyte monolayer of the neonatal extrahepatic bile duct was immature, lacking the uniform apical glycocalyx and mature cell-cell junctions typical of adult cholangiocytes. Functional studies showed that the glycocalyx protected against bile acid injury and that neonatal cholangiocyte monolayers were more permeable than adult monolayers. In adult ducts, the submucosal space was filled with collagen I, elastin, hyaluronic acid, and proteoglycans. In contrast, the neonatal submucosa had little collagen I and elastin, although both increased rapidly after birth. In vitro modeling suggested that the composition of the neonatal submucosa relative to the adult submucosa led to increased diffusion of bile. A Col-GFP reporter mouse showed that cells in the neonatal but not adult submucosa were actively producing collagen.

**Conclusion:** We identified four key differences between the neonatal and adult extrahepatic bile duct. We showed that these features may have functional implications, suggesting the neonatal extrahepatic bile ducts are particularly susceptible to injury and fibrosis.

**Lay Summary:** Biliary atresia is a disease that affects newborns and is characterized by extrahepatic bile duct injury and obstruction with resulting liver injury. We identify four key differences between the epithelial and submucosal layers of the neonatal and adult extrahepatic bile duct and show that these may render the neonatal duct particularly susceptible to injury.

**Highlights:**

1. The apical glycocalyx is thin and patchy in neonatal compared to adult cholangiocytes
2. Neonatal cholangiocytes have immature cell-cell junctions and increased permeability
3. The neonatal submucosal space has minimal collagen I or elastin
4. The neonatal submucosal space contains many actively collagen-secreting cells

**Graphical abstract:** 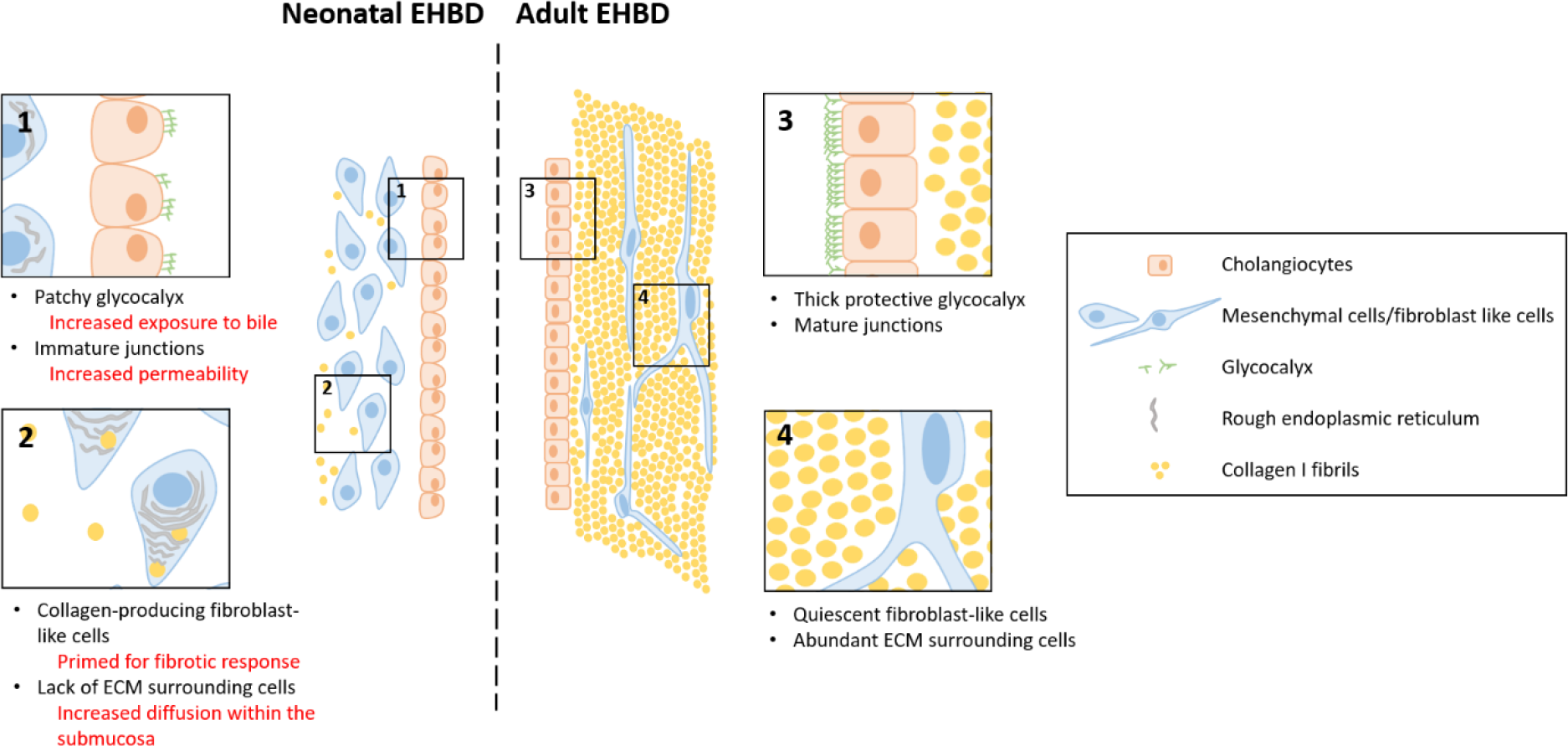

## Introduction

Mammalian bile travels through an arborizing network encompassing bile canaliculi, the canals of Hering and ductules, and cholangiocyte-lined intrahepatic (IHBD) and extrahepatic (EHBD) bile ducts. The IHBD and EHBD arise from different primordia: the cranial portion of the endodermal diverticulum gives rise to the liver and the IHBD while the caudal portion develops into the EHBD (1–3). The EHBD and IHBD, in addition to differing in development, size, and surrounding tissue, are also notable for differential susceptibility to certain cholangiopathies including primary biliary cholangitis, primary sclerosing cholangitis, and the neonatal fibro-obliterative disease biliary atresia (BA). BA, which at the time of presentation primarily affects the EHBD, is specific to neonates, who are born apparently healthy but progress remarkably quickly (within several months) to liver fibrosis and cirrhosis. Recent human and animal model data suggest that the initial insult is prenatal, affecting the neonate but sparing the mother (4–8). This suggests a paradigm whereby the fetal and neonatal ducts, especially the EHBD, are uniquely susceptible to initial and progressive damage from injurious agents.

Bile ducts have specialized anatomical features that protect them from bile acid-mediated injury and other insults. Specialized protective features of cholangiocytes include unique apical membranes and an apical glycocalyx with a protective “bicarbonate umbrella” layer (9–11). The EHBD submucosa may also have a protective function. It has recently been described to comprise a fluid-containing interstitial space with a lattice of collagen bundles, elastin, and thin fibroblast-like cells directly adherent to the collagen bundles (12). These features serve a barrier function, provide structural support to the duct, and also likely serve as a shock absorber and a conduit for the movement of soluble and cellular factors (12).

The development of these potential protective features in the neonatal EHBD has not been reported. Similarly, the mechanism of specific neonatal and EHBD susceptibility to injury is unknown, although we and colleagues previously reported that the EHBD has low levels of glutathione compared to the IHBD and liver (13, 14). Here we report a detailed anatomical and functional study of the murine EHBD, including mucosal and submucosal layers, from birth through adulthood. We describe the identification of four potentially developmentally-coordinated features that may in part explain the particular susceptibility of the neonatal EHBD to injury and fibrosis. We focused on BALB/c mice; this strain, unlike C57BL/6 and other common laboratory strains, is sensitive to neonatal infection with rhesus rotavirus and is used to generate the widely-used rhesus rotavirus infection model of BA (8).

## Methods

### Chemicals

Reagents were obtained from Sigma-Aldrich (St. Louis, MO, USA), unless otherwise noted.

### Animals

All work with mice was in accordance with protocols approved by the University of Pennsylvania and University of California, San Diego Institutional Animal Care and Use Committees, as per the National Institutes of Health Guide for the Use and Care of Animals. BALB/c mice were obtained from the Jackson Laboratory (Bar Harbor, ME, USA). Collagen α1(I)-GFP (Col-GFP) mice were as previously described (15, 16). Both male and female mice were used for all analyses. Pig bile was a gift from Robert Mauck (University of Pennsylvania, PA, USA) and was obtained from animals previously euthanized as part of other studies.

### Human samples

Anonymized human EHBD sections were obtained from the Anatomic Pathology archives of the Children’s Hospital of Philadelphia, with IRB approval. Formalin-fixed paraffin-embedded EHBD sections were obtained from fetuses stillborn at 22, 34, 36 and 39 weeks gestation during the course of autopsies performed with parental consent, and from a native liver explant from a 7 year-old child and a partial hepatectomy specimen resected for a primary pancreatic tumor from a 10 year-old.

### Histology, immunostaining, and lectin staining

EHBDs and livers were dissected from mice at the ages noted and were either formalin fixed and paraffin embedded or frozen in O.C.T. (Tissue Tek, VWR, Bridgeport, NJ, USA), then sectioned at a 5 µm thickness. For antibody staining, the antibodies used are listed in Supplemental Tables 1 and 2. For lectin staining, slides were washed briefly in PBS and stained using biotinylated lectins per the manufacturer’s protocol (Vector Laboratories, Burlingame, CA, USA). The lectins used are shown in Supplemental Table 3. For mucopolysaccharide detection, tissues were stained using Alcian blue. Nuclei were stained with 4′,6-diamidino-2-phenylindole (DAPI).

Further details on staining are found under Supplemental Methods.

### Transmission electron microscopy (TEM)

EHBDs and livers from neonatal and adult mice were fixed (2.5% glutaraldehyde, 2% paraformaldehyde, 0.1M sodium cacodylate buffer, pH 7.4) in situ immediately after euthanasia and then dissected and fixed in the same buffer overnight at 4°C. After subsequent buffer washes, the samples were post-fixed in 2.0% osmium tetroxide for 1 h at room temperature and rinsed in dH_2_O prior to en bloc staining with 2% uranyl acetate. After dehydration through a graded ethanol series, the tissue was infiltrated and embedded in EMbed-812 (Electron Microscopy Sciences, Fort Washington, PA, USA). Thin sections were stained with uranyl acetate and lead citrate and examined with a JEOL 1010 electron microscope (JEOL USA, Peabody, MA, USA) fitted with a Hamamatsu digital camera (Bridgewater, NJ, USA) and AMT Advantage image capture software (Advanced Microscopy Techniques Corp., Woburn, MA, USA). Ruthenium red staining of the sections was performed as described (11).

### Image analysis

Image analysis was performed using ImageJ. The thickness of the glycocalyx was measured in TEM images; 10 random regions were selected from 3 independently isolated EHBDs per age group. Measurement of collagen diameter in TEM images was done using the threshold and particle analysis functions of ImageJ. For each age group, 150-300 fibrils were counted for 3-5 independent EHBDs. Quantification of immunohistochemistry and second harmonic generation (SHG) signal was done using the threshold and % area functions.

### Primary cholangiocyte isolation and culture

Primary extrahepatic biliary epithelial cells were isolated from adult and neonatal mice, cultured in BEC media as previously described (17), and used up to passage 3.

### Flow cytometry

Primary adult cholangiocytes were cultured to 80% confluence on a 10 cm plate on collagen at 37°C. Cells were trypsinized, suspended in serum-free BEC medium and incubated with or without 0.1 U/ml (Sigma) neuraminidase in serum-free BEC medium for 2.5 h. Cells were washed with BEC medium containing 10% FBS and were re-suspended in Flow Cytometry Staining Buffer (1X) (R&D Systems, Minneapolis, MN, USA) supplemented with 1% FBS. They were incubated with 1 µg/ml of FITC-soybean agglutinin (SBA) lectin (Vector Laboratories, Burlingame, CA, USA) for 30 min. Cells were then washed and resuspended in FACS buffer and analyzed for lectin binding affinity by performing flow cytometry analysis on an LSR II equipped with FACS DiVA software (version 6.1.1; BD). Measurement in the absence of lectins was performed as a negative control and at least 1 × 10^4^ cells were analyzed for each sample. Data were analyzed using FlowJo software (version 10; FLOWJO, LLC). The data were manually gated in FlowJo and then Flow SOM was performed using the plugin installed in the software (18). A minimal spanning tree was constructed to visualize the expression of FITC.

### Cell viability assays

Confluent neonatal and adult cholangiocytes were cultured on thin collagen cells at 1 mg/ml. They were incubated with or without 0.1 U/ml of neuraminidase in serum-free BEC medium for 2.5 h. The medium was then replaced with BEC medium containing 10% FBS and bile acids or diluents and the cells were treated for 4 h. Stock solutions of chenodeoxycholic acid (CDC) were made in ethanol/DMSO (50:50, v/v); glycochenodeoxycholic acid (GCDC) stocks were made in HBSS, both at pH 6.8-7.0. The bile acids were diluted in BEC medium at 1 mM for GCDC and 0.5 mM for CDC (11). Cell viability was assessed by the WST-1 assay (Sigma-Aldrich) following the manufacturer’s protocol.

### Spheroid culture and rhodamine efflux assay

Primary neonatal and adult mouse extrahepatic cholangiocytes were cultured in 3D in a collagen-Matrigel mixture for 7 days as described previously and a rhodamine efflux assay was then performed (13). Briefly, cells were incubated in Rhodamine 123 for 15 min to load the lumens; live cells were then washed and imaged every 20 minutes using a spinning disk confocal microscope with a wide 40X lens. Corrected total cell fluorescence (CTCF) was measured using ImageJ (v1.48, NIH) as previously described (19). The percentage decrease of fluorescence was calculated using the initial 20 min time point as the reference point.

### Second harmonic generation imaging

Fixed whole EHBDs were imaged using a Leica SP8 confocal/multiphoton microscope and Coherent Chameleon Vision II Ti:Sapphire laser (Leica, Buffalo Grove, IL, USA) tuned to a wavelength of 910 nm. SHG of collagen generates both forward and backward signal; directionality of scatter is dependent on the size and orientation of the fibril, with backward scatter associated with immature fibrils (20).

### In vitro bile diffusion assay

We made an in vitro model of the submucosal matrix by mixing rat tail collagen I (Corning, NY, USA) at a concentration of 0.25, 0.5, 1, 2, and 5 mg/ml with high molecular weight (15 MDa) hyaluronic acid (HA) (Lifecore Biomedical, Chaska, MN, USA) at a concentration of 0.5 mg/ml in a buffered medium (HEPES, NaCO_3_, NaOH, and DMEM). A channel with a diameter of 160 μm was made in the gel mixture by inserting an acupuncture needle before gelation. 50 μl of Yucatan pig bile (which is autofluorescent) was flowed through the channel and the gel mixture was imaged every 5 seconds for 2 min using an EVOS FL Auto imaging system. The diffusion rate (D^2^/(2xT), where D = distance, and T = time) over the first 30 secs was calculated for 10-12 gels per collagen concentration.

### Statistical analysis

Statistical significance was assessed using one-way ANOVA followed by Tukey’s post hoc analysis. p<0.05 was regarded as statistically significant, and calculated with Prism 7 (GraphPad Software, La Jolla, CA, USA). An unpaired t test was used to compare spheroid sizes, with P values <0.05 considered to represent statistical significance.

## Results

### Neonatal BALB/c mouse cholangiocytes lack an apical glycocalyx

We first assessed whether neonatal mouse cholangiocytes have an apical glycocalyx, as has been reported for adult human and mouse cholangiocytes (10, 21). TEM images of livers and EHBDs from BALB/c mice at multiple timepoints from postnatal day 0 through adulthood confirmed published data that adults have an apical glycocalyx (Fig. 1) (10, 11). The glycocalyx of both the EHBD and IHBD of neonates (days 0-2), however, was thin and sparse (Fig. 1 A, B and Supplemental Fig. 1). The glycocalyx thickness of newborn bile ducts ranged from 0-10 nm; the glycocalyx layer was continuous and increased in thickness to about 20 nm by postnatal day 7 and 25-30 nm in adults (Fig. 1B). (Note that fixation dehydrates the glycocalyx and it is likely to be significantly thicker in vivo; the relative thickness of different samples, however, holds true.) Although the glycocalyx was easily visualized by TEM even when unstained, its identity was confirmed by staining with ruthenium red and osmium tetroxide, which bind to glycosaminoglycans present in the glycocalyx (Fig. 1C) (22, 23). EHBDs were also stained with Alcian blue, which indicated the presence of acidic glycans on the apical surface of adult but not neonatal EHBDs (Fig. 1D). Thus, BALB/c mice had a poorly developed IHBD and EHBD glycocalyx at birth, although it became continuous and thicker during the first two weeks of life.

**Fig. 1.**
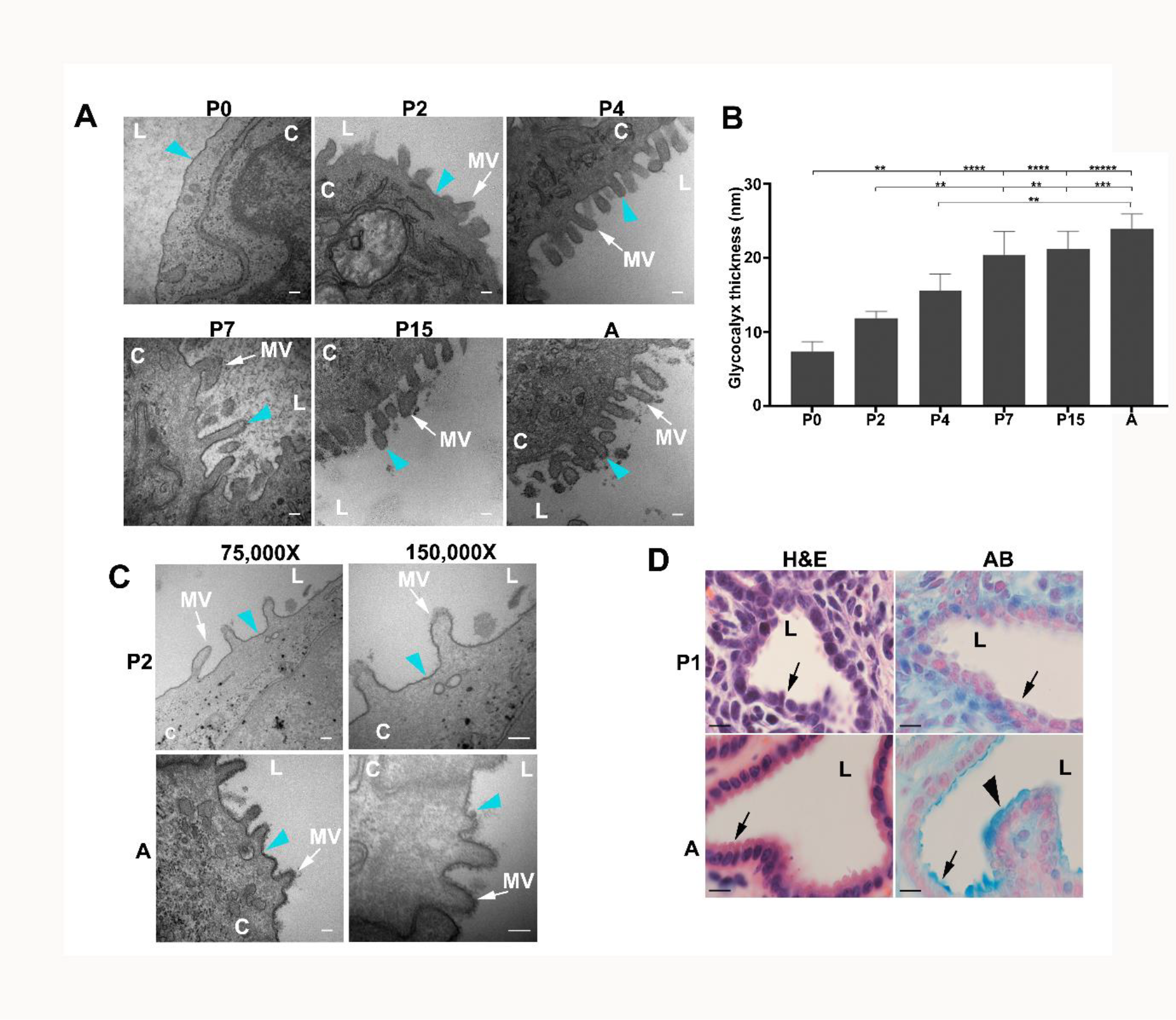
BALB/c neonatal EHBDs have a patchy glycocalyx. (A) TEM images of EHBDs from BALB/c-mice at postnatal days 0 (P0), 2 (P2), 4 (P4), 7 (P7), 15 (P15) and adult (A). Teal arrowheads point to the glycocalyx on microvilli (MV; white arrows). Scale bars, 100 nm (B) Thickness of the glycocalyx layer from TEM images was measured on 3 EHBDs per age group. Data shown are mean ± SD, ** P<0.01, *** P<0.001, **** P<0.0001. (C) TEM images of an adult (A) and postnatal day 2 (P2) EHBD after Ruthenium red staining. Teal arrowheads show the glycan coat stained in black; white arrows point to microvilli (MV). Scale bar is 100 nm. (D) Alcian blue (AB) staining of adult (A) and postnatal day 1 (P1) EHBD with corresponding H&E images; mucins in the glycocalyx appear blue. Images shown are representative of staining from 3 EHBDs per age group. Black arrows point to the cholangiocyte layer while bold black arrowheads point to the mucin stained glycocalyx. Scale bars, 10 µm. **L** denotes lumen and **C** cholangiocytes.

### The cholangiocyte glycocalyx varies during development and protects cells from bile toxicity

To understand the complexity of the glycocalyx, we stained neonatal and adult BALB/c mouse EHBD sections using a panel of lectins (Supplemental Table 3). Lectins *Concanavalin A* (Con A), *Phaseolus vulgaris Erythroagglutinin* (PHA-E), *Phaseolus vulgaris Leucoagglutinin* (PHA-L), *Peanut Agglutinin* (PNA), *Griffonia (Bandeiraea) simplicifolia* (GSL), and *Soybean* (SBA), out of a panel of 12, showed strong binding to the EHBD glycocalyx of adult BALB/c mice (Fig. 2). The apical surface of neonatal EHBDs, however, remained unstained with all except the PHA-E and SBA lectins, which showed faint discontinuous staining. Cholangiocytes at postnatal day 7 had light staining with PHA-E, PHA-L, and ConA, as well as staining with the lectin *Maackia amurensis II* (MAlII), which binds only carbohydrate structures that contain sialic acid in an (α-2,3) linkage; MalII did not stain postnatal day 1 EHBDs or adult bile ducts (Supplemental Fig. 2). Lectin staining of well-preserved human EHBDs was consistent with results from mice. PHA-E staining showed a glycocalyx on the luminal surface of EHBDs from 7 and 10 year old patients, while there was no PHA-E staining of EHBDs taken from stillborn infants at gestational ages up to 39 weeks (Supplemental Fig. 3). These human samples appeared well preserved, but degradation of the glycocalyx in utero cannot be ruled out. Overall, based on lectin staining, the glycocalyx develops and remodels after the ducts themselves form.

**Fig. 2.**
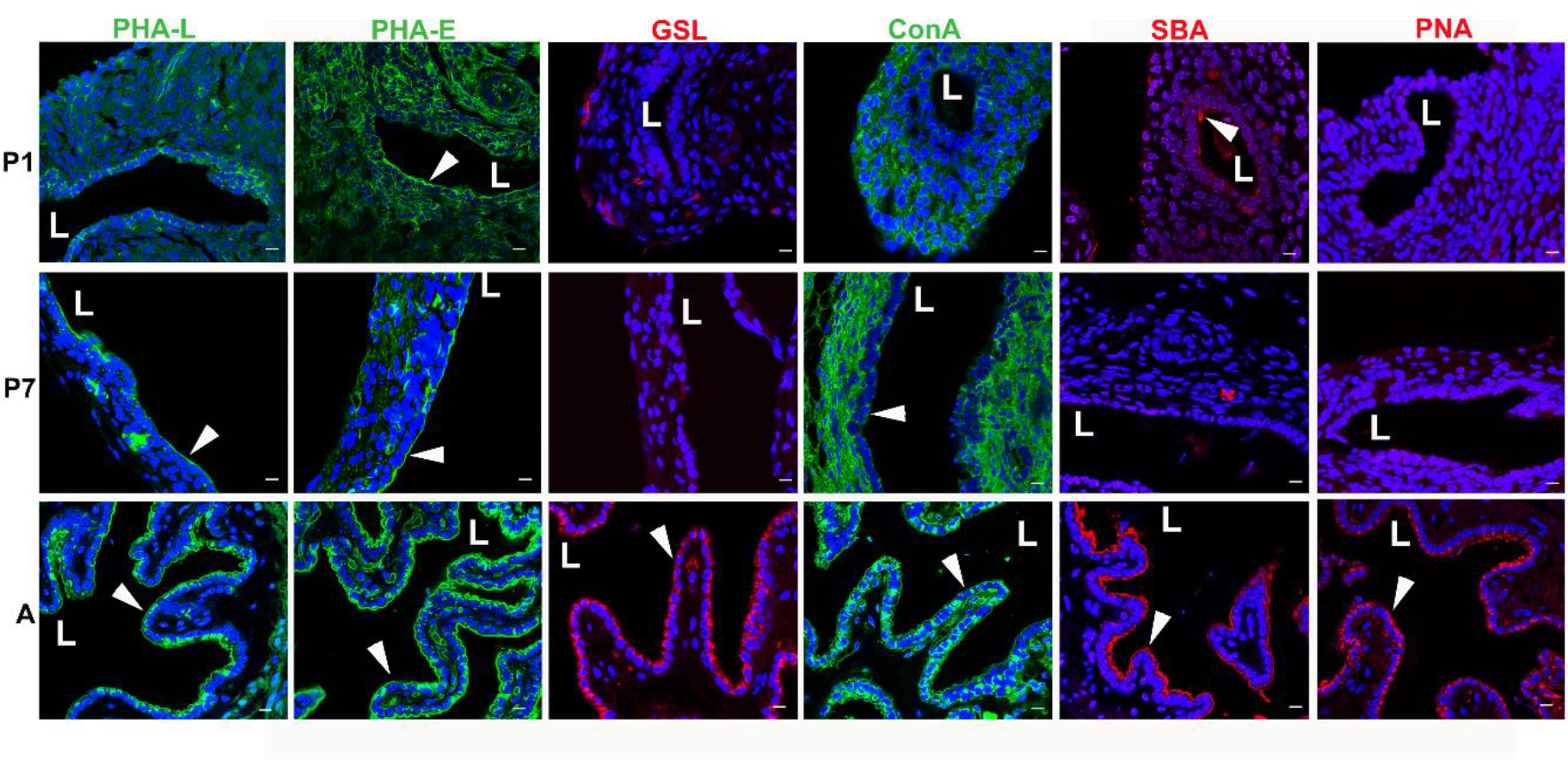
Lectin staining of the glycocalyx in BALB/c mice. EHDBs from BALB/c mice at postnatal days 1 (P1) and 7 (P7) and in adults (A) were stained using a lectin panel as per Supplemental Table 3. White arrowheads point to the glycocalyx on the apical side of the cholangiocyte monolayer facing the duct lumen (L). Lectin staining is red or green; nuclei were stained with DAPI (blue). Representative images shown of EHBDs from 3 animals stained per age group. Scale bars, 10 μm.

**Fig 3.**
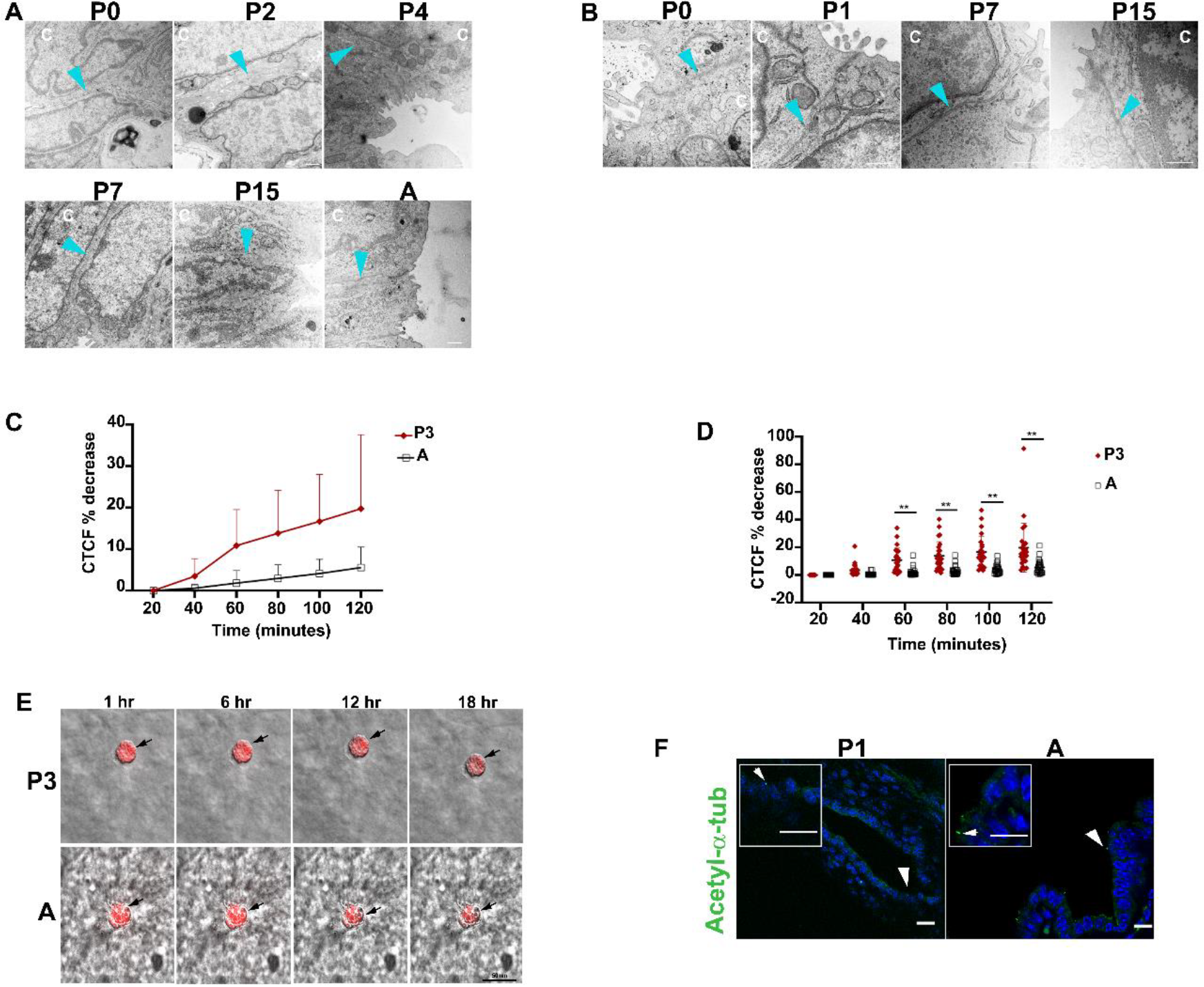
Cholangiocyte cell-cell junctions are immature in neonatal mouse EHBDs. TEM images of (A) EHBDs and (B) IHBDs from BALB/c-mice at different postnatal ages: day 0 (P0), 1 (P1), 2 (P2), 4 (P4), 7 (P7), 15 (P15) and adult (A), magnification X25,000 (A) and X50,000 (B). Teal arrowheads point to cell-cell junctions between cholangiocytes. Representative images shown from 3 ducts examined per age group. Scale bars, 500 nm. C,D) Cholangiocytes spheroids were loaded with Rhodamine 123 and the loss of rhodamine (CTCF % decrease) was measured over 2 hrs (n=25 P3, 26 adult). The plots show % decrease in fluorescence. Data shown are mean ± SD * P<0.05. E) Black arrows point to the accumulated rhodamine (red) in the lumens of typical spheroids. Scale bar, 50 μm. F) EHBDs from postnatal day 1 (P1) and adult (A) BALB/c mice were stained for acetylated α-tubulin, a marker of primary cilia. Nuclei are stained with DAPI (blue). White arrows point to examples of stained cilia. Representative images shown of EHBDs from 3 animals stained per age group. Scale bars, 10 μm. Scale bars of inset boxes, 20 μm.

In order to determine whether the glycocalyx plays a protective role in mice, as reported previously for adult human cholangiocytes (11), we isolated primary cholangiocytes from adult mouse EHBDs (Supplemental Fig. 4A), then enzymatically treated them with neuraminidase, which cleaves negatively-charged terminal sialic acids in the glycocalyx (10, 11). The efficacy of the neuraminidase treatment was confirmed by flow cytometry analysis using adult cholangiocytes labeled with the lectin SBA, which binds to newly exposed sugar residues in the glycocalyx after sialic acid removal (Supplemental Fig. 4B) (10). Neuraminidase-modified adult cholangiocytes were then exposed to the bile salts CDC and GCDC and toxicity was analyzed using a WST 1 assay. Both CDC and GCDC are known to be toxic because of high pKas (11). We observed that both these bile acids were toxic to primary cholangiocytes at pH values below 7; however, disruption of the glycocalyx resulted in increased susceptibility to bile acid-mediated injury (Supplemental Fig. 4C).

**Fig 4.**
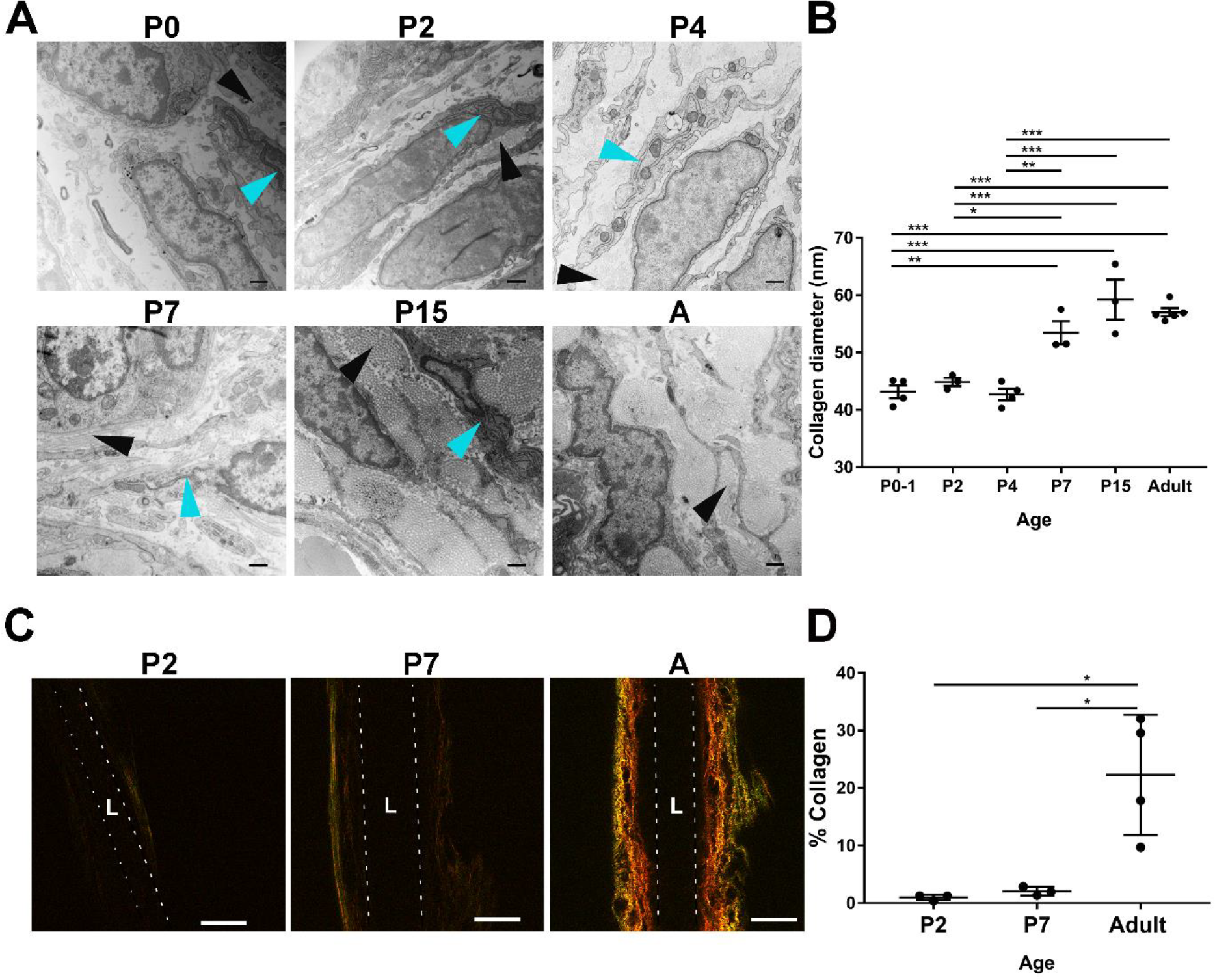
Collagen is deposited after birth in the mouse EHBD submucosa. (A) TEM images from mouse EHBDs isolated from BALB/c mice at postnatal days 0 (P0), 2 (P2), 4 (P4), 7 (P7), and 15 (P15) and adults (A). Scale bars, 500 nm. Black arrowheads show collagen fibrils; teal arrowheads show rough endoplasmic reticulum in cell cytoplasm. (B) Collagen fibril diameters were measured using Image J. For each age group, 150-300 fibrils were counted for 3-5 independent EHBDs. Data shown are mean ± SEM, * P<0.05, ** P<0.01, *** P<0.001. (C) Second harmonic generation imaging of whole ducts to visualize collagen, with all imaging carried out at the same settings. Forward signal is seen in red and backward signal in green. Dotted lines outline the lumen (**L**) walls. Scale bars, 100 μm. (D) Quantification of combined forward and backward SHG signal of 3 EHBDs per age group. Data shown are mean ± SD, * P<0.05.

### Cell-cell junctions of neonatal EHBD, not IHBD, are immature

Because cholangiocyte cell-cell junctions are key to protecting the basolateral surfaces of cholangiocytes and the submucosa from bile acid toxicity, we compared these junctions in mice of different ages. On TEM imaging, junctions in BALB/c mouse EHBDs from birth through postnatal day 7 appeared immature, with poorly developed tight junctions and, in many cases, open channels from apical surface to basement membrane (Fig. 3A). No significant differences were observed between adult cholangiocytes and cholangiocytes of the IHBD, even at birth (Fig. 3B).

To determine the functional relevance of these differences, we measured leakage of Rhodamine 123 from adult and neonatal mouse cholangiocyte spheroids in 3D culture. Rhodamine 123 is a fluorescent dye taken up into cholangiocytes via the basal surface which is then effluxed into the lumen by the apical transporter MDR1 and (to a lesser extent) MRP1. In studies with adult cholangiocyte spheroids, leakage of rhodamine from the lumen is minimal (13, 24, 25). Spheroids with well-formed lumens were loaded for 15 minutes (13), then imaged every 20 minutes to assess leakage (Fig. 3E). Neonatal and adult spheroids were of similar diameters (Supplemental Fig. 5), although total rhodamine uptake into neonatal lumens was about half that for adult lumens. Neonatal cholangiocytes demonstrated a rapid initial loss of signal, consistent with the anatomical differences observed in their junctions (Fig. 3C, D). This may be related to developmental variability in the transporters mediating uptake; nevertheless, our data strongly suggest that neonatal EHBD cholangiocyte monolayers are more permeable that adult EHBD cholangiocyte monolayers.

**Fig 5.**
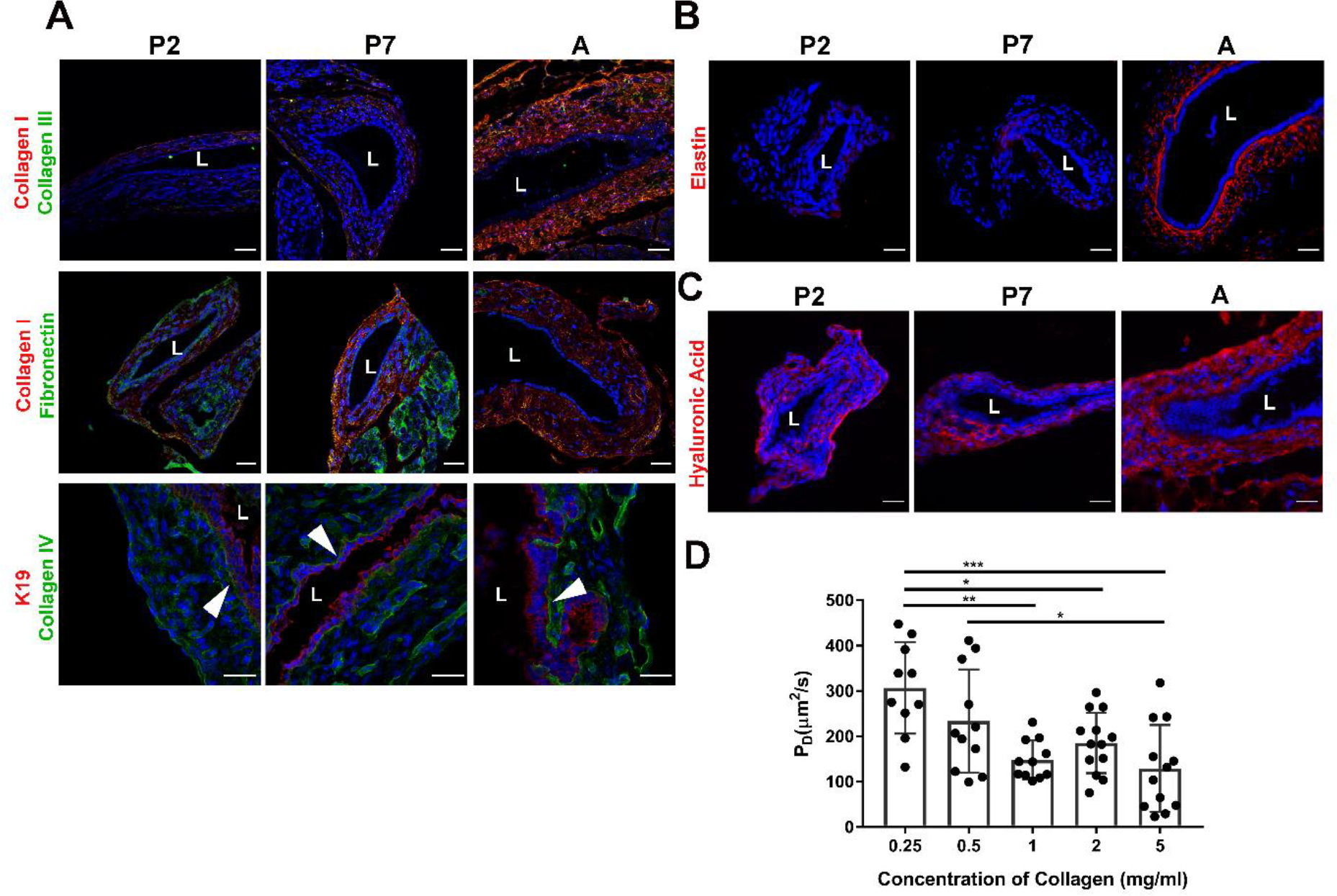
Expression of ECM proteins varies in the neonatal and adult mouse EHBD. (A) EHBDs from BALB/c mice at postnatal days 2 (P2) and 7 (P7) and adults (A) were stained for collagen I (red) or the cholangiocyte marker K19 (red) and either collagen III (green), fibronectin (green), or collagen 4 (green). White arrowheads (bottom panel) show the basement membrane of cholangiocytes. (B,C) EHBDs were also stained for (B) elastin (red) and (C) hyaluronic acid (red). Nuclei are stained with DAPI (blue). Representative images shown from 3-4 EHBD stained per age group and condition. Scale bars, 30 μm. **L** denotes the lumen. (D) Diffusion rate of bile from cylindrical lumens in HA (0.5 mg/ml) and collagen gels (0.25-5 mg/ml). A total of 10-12 gels were tested per condition. Data shown are mean ± SD, * P<0.05, ** P<0.01, *** P<0.001.

Although cell-cell junctions were immature, other cholangiocyte structures appeared mature even in neonates. Apical microvilli were abundant on EHBD and IHBD cholangiocytes from mice at all ages, and acetylated-α-tubulin staining was positive for cilia in neonates and adult EHBDs (Fig. 3F and Supplemental Fig. 6).

**Fig 6.**
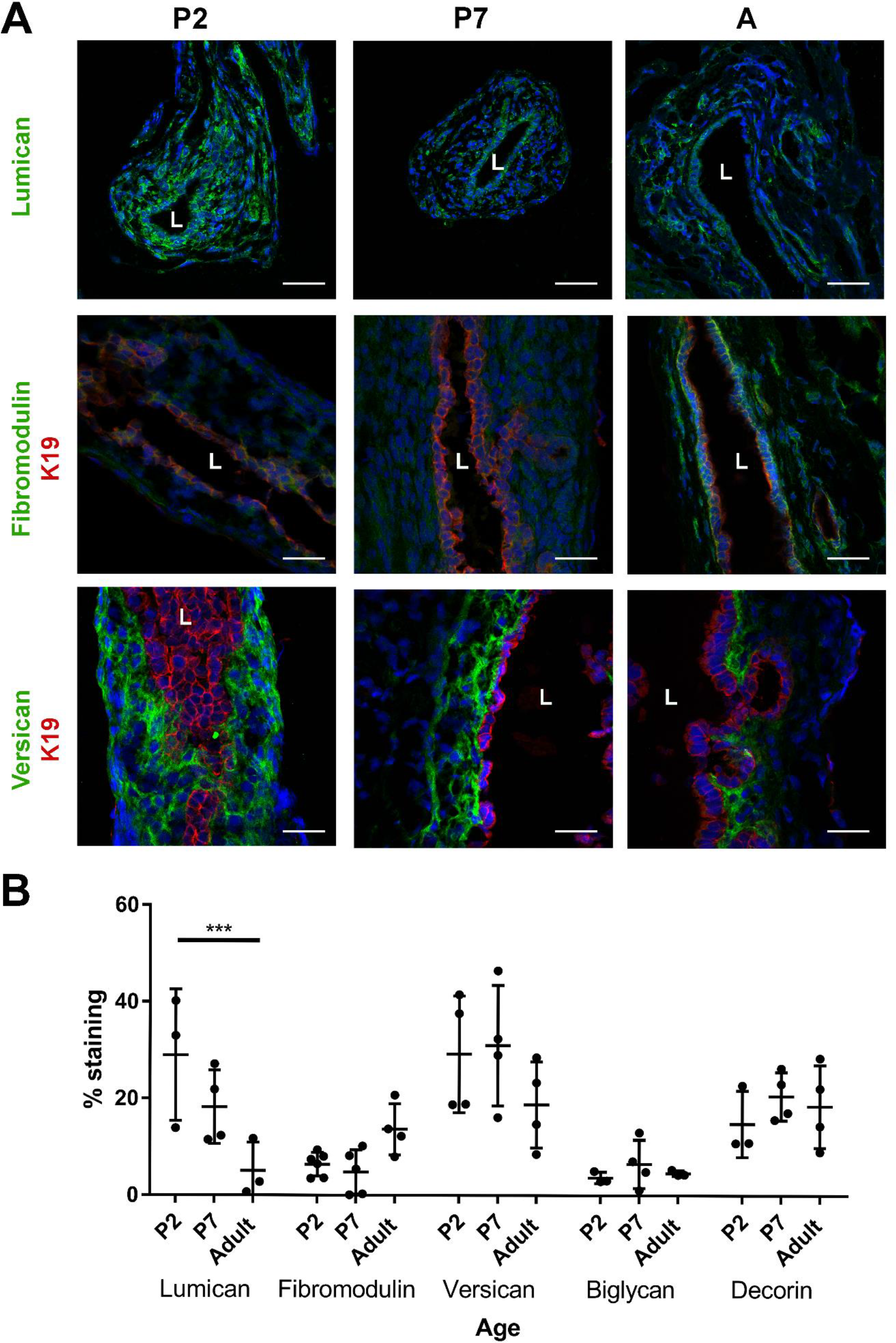
Presence and distribution of proteoglycans in the EHBD submucosa varies with age. EHBDs from BALB/c mice at postnatal days 0 (P0), 1 (P1), and 7 (P7) and adults (A) were stained with antibodies against collagen (red) and the proteoglycans lumican, fibromodulin and versican (all green). Nuclei were stained with DAPI (blue). Scale bars, 30 μm. **L** denotes the lumen. (B) Quantification of proteoglycan staining on 3-6 EHBD per age group. Data shown are mean ± SD, *** P<0.001.

### Collagen is deposited in the EHBD submucosa in the postnatal period

We examined the architecture of the mouse EHBD submucosa as a function of age. The adult mouse EHBD submucosa, similar to the submucosa of adult human ducts (12), contained a network of collagen bundles. In contrast, the neonatal (postnatal days 1-4) duct submucosa had only scattered collagen fibers (Fig. 4A, black arrows); the space appeared mostly empty on TEM, and immunostaining showed that it was filled with ground substance (HA and proteoglycans; Figs. 5C & 6). In neonatal ducts (postnatal day 2), collagen contributed to less than 1% of the duct area, however, in adults this increased to over 20% (Fig. 4D). Collagen fibril diameter significantly increased from 45 nm to 55 nm between postnatal days 4 and 7 (Fig. 4B), after which the diameter remained the same through adulthood. EHBDs of all ages were assessed for the presence of fibripositors (26), structures reported to mediate collagen alignment under tension; however, few fibripositors were observed in the EHBD, and these were only in the neonatal EHBD (Supplemental Fig. 7). Collagen was deposited first in the outer layer of the duct wall, as shown by SHG microscopy as well as immunostaining for the fibrillar collagens I and III (Figs. 4C, 5A).

**Fig. 7.**
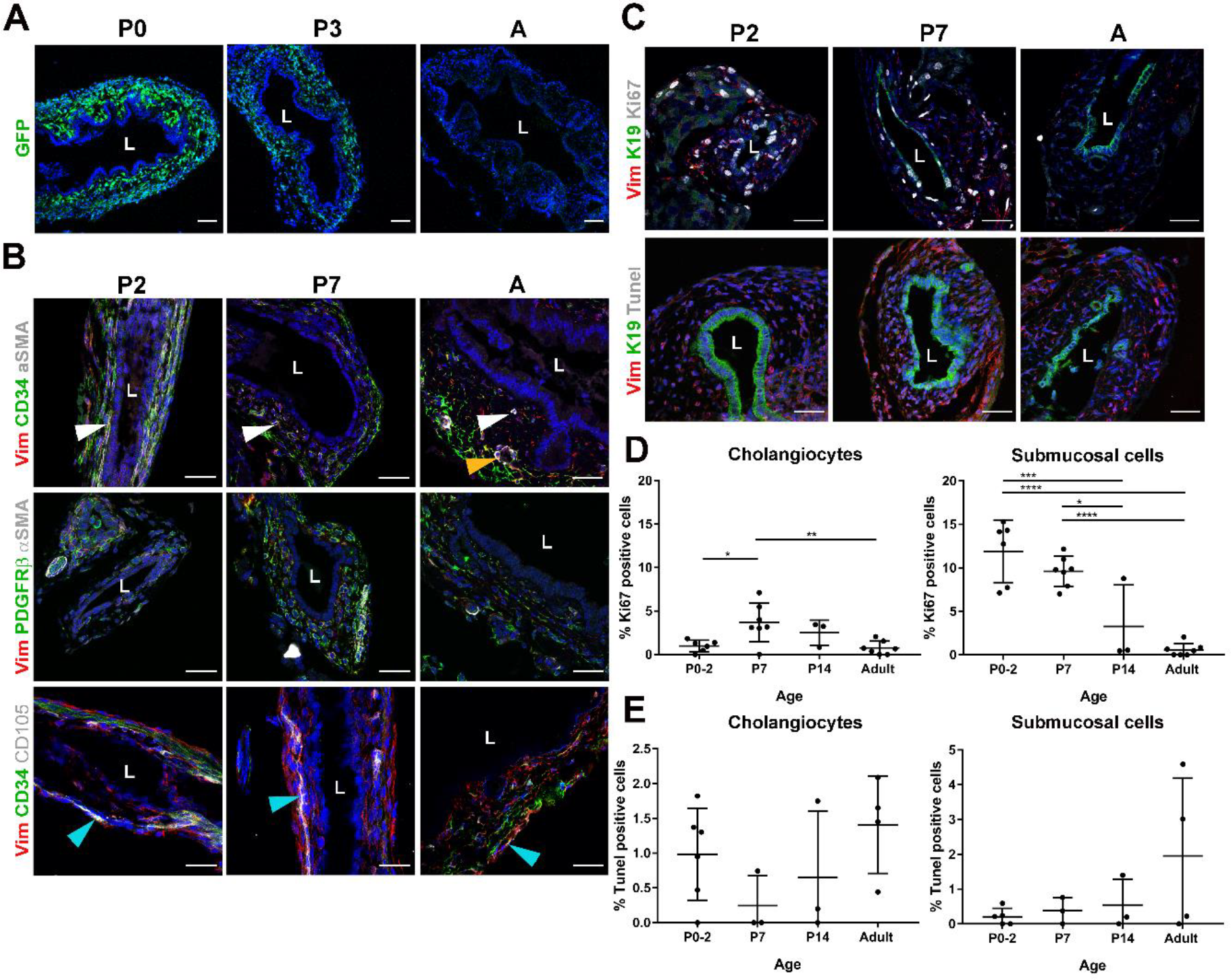
A heterogeneous population of cells is present in the developing EHBD. (A) EHBDs from Col-GFP mice at postnatal days 0 (P0) and 3 (P3) and adults (A) were immunostained for GFP. (B) EHBDs isolated from BALB/c mice at postnatal days 2 (P2) and 7 (P7) and adults (A) were immunostained with antibodies against the mesenchymal marker vimentin (red) and either the endothelial marker CD34 (green), or fibroblast markers PDGFRβ (green) and αSMA (grey) or the mesenchymal stem cell marker CD105 (gray). White arrowheads show cells positive for αSMA, CD105, Vimentin and CD34. Orange arrowhead shows αSMA-positive blood vessel. Teal arrowheads show cells positive for CD105, Vimentin, and CD34. (C) EHBDs were also stained for vimentin (red) for submucosal cells and K19 (green) for cholangiocytes and either Ki67 (gray) for proliferation or TUNEL staining (gray) for cell apoptosis. Nuclei were stained with DAPI (blue). Representative images from 3-4 independent EHBD stained per age group. (D) Ki67 and (E) TUNEL staining were quantified separately for the cholangiocyte monolayer and for the submucosa on 3-4 EHBD per age group. Data shown are mean ± SD, *** P<0.001, **** P<0.0001. Scale bars, 30 μm. **L** denotes the lumen.

To model the effect of increasing collagen deposition on interstitial flow through the EHBD submucosa, mixtures of high molecular weight HA (0.5 mg/ml) and increasing collagen I concentrations (0.25-5 mg/ml) were gelled around a needle to form a channel and the diffusion rate of bile from the channel into the mixture was measured (Fig. 5D). Diffusion rates were inversely correlated with collagen concentration, suggesting that bile leakage into the submucosa may be limited more by collagen in the adult than in the neonatal EHBD.

### ECM composition in the EHBD submucosa varies with age

The composition of the ECM in the EHBD submucosa changed markedly with age. In adults, the submucosa contained collagens I, III (small amounts) and IV, fibronectin, HA and the proteoglycans lumican, decorin, fibromodulin and versican, but not the proteoglycans biglycan and aggrecan (Figs. 5A-C, 6, and Supplemental Fig. 8). Collagen IV was found at the edge of the outer wall, as well as along the basement membrane (with a thick layer of elastin), while versican deposition was concentrated around the submucosal cells under the cholangiocyte layer (Fig. 5A). In contrast, the neonatal duct had no or little collagen I, III, or elastin (Fig. 5A, B), suggesting decreased tissue integrity and ability to respond to mechanical stress. The neonatal submucosa did, however, contain proteoglycans (although not fibromodulin) and HA (Fig. 5C, 6). Notably, lumican, which plays a role in collagen fibrillogenesis, was more highly expressed in neonatal EHBDs (Fig. 6B). Lectin staining of the submucosa with PHA-L, ConA and PHA-E showed that specific glycosaminoglycan epitopes change with age (Fig. 2). Thus, the amount and composition of the extracellular matrix in the submucosal space changed markedly with age, suggesting the potential for significant differences in submucosal fluid flow, cell movement and mechanical properties including the response to obstruction.

### Neonatal submucosal cells are fibrogenic

TEM images showed that the submucosal cells of the EHBD varied in morphology with age (Fig. 4A). In adult mouse ducts, most submucosal cells were flat, with long thin processes that surrounded and directly adhered to adjacent collagen bundles, as has been reported for human ducts (Fig. 4A and (12)). These cells had little rough endoplasmic reticulum (RER). In neonates, however, most submucosal cells were round and contained large amounts of RER, suggesting high synthetic activity. The presence of actively collagen-secreting cells in neonates but not adults was confirmed by using neonatal Col-GFP reporter mice, which showed significantly more GFP expression in neonates than in adults (Fig. 7A). The submucosal cells in the developing neonatal duct were highly proliferative, as shown by Ki67 staining (Fig. 7C, D). TUNEL staining demonstrated that these cells were also less apoptotic, suggesting that turnover of cells during postnatal development of the EHBD was minimal (Fig. 7C, E).

In EHBD of all ages, the majority of cells stained positively for the mesenchymal and fibroblast cell markers vimentin, CD34 and PDGFRβ (Fig. 7B). In adults, the expression pattern of CD34 varied with the relative position of the cells within the submucosa, with expression at its highest in the outer layer of the duct. α-SMA staining was observed, as expected given the Col-GFP data, in the smooth muscle cells surrounding vessels in both the neonate and adult; however, a subpopulation of the CD34-and vimentin-positive cells in the neonatal submucosa also stained for α-SMA. A second subpopulation of submucosal cells, present most prominently at postnatal day 7, expressed the mesenchymal stem cell marker CD105. No fat droplets were seen in any fibroblast-like cells on TEM analysis, suggesting that they are not retinoid-storing like hepatic or pancreatic stellate cells. The submucosa of the EHBD was also interspersed with F4/80 positive macrophages and was crossed by nerves (Supplemental Figs. 9 and 10).

## Discussion

We report here a systematic study of the postnatal development of the murine EHBD mucosa and submucosa that has implications for understanding why neonatal ducts are uniquely targeted for injury in BA. We identify four major differences between neonate and adult: 1) neonatal cholangiocytes lack a protective apical glycocalyx; 2) neonatal EHBD tight junctions in the cholangiocyte monolayer are immature and cholangiocytes from neonates are leakier than from adults; 3) the neonatal submucosa lacks the major structural proteins collagen I and elastin, potentially impacting diffusion of bile leaked into the submucosal space as well as the mechanical properties of the duct; and 4) the neonatal submucosa contains proliferative cells that are actively secreting collagen even in the absence of injury. Collectively, these suggest that the neonatal EHBD has increased susceptibility to injury and fibrosis.

The apical glycocalyx is thought to incorporate a “bicarbonate umbrella” and to serve as an important defense mechanism protecting cholangiocytes from bile acids, particularly those like glycochenodeoxycholic acid (GCDC) with high pK_a_ values (10, 21). Published work shows that human and mouse adult cholangiocytes express a sialated apical glycocalyx with a high concentration of α1-2 fucose links, and that bile acid-induced cholangiocyte damage is worse when this glycocalyx is disrupted with agents such as neuraminidase (10). Our TEM and lectin staining shows that this glycocalyx layer is thin and discontinous in BALB/c mice in the first few days of life, and that it is similarly absent before birth in the human EHBD.

Characterization of the glycocalyx via lectin binding offers insight into the composition and abundance of certain sugars and highlights age-related differences in the glycoconjugate composition of the glycocalyx (17, 27). The PNA lectin, which recognizes only unsialylated galactose residues (28), bound to the EHBD glycocalyx only in adults. The SNA lectin, which binds to sialic acid attached to terminal galactoses via α-2,6 and to a lesser degree, α-2,3 linkages, strongly stained the glycocalyx of adult cholangiocytes while SBA, which preferentially binds to oligosaccharide structures with terminal α- or β-linked N-acetylgalactosamines, stained both adult and neonatal glycocalyces.

To investigate the functional relevance of the glycocalyx in cholangiocytes, we performed bile acid toxicity studies with or without enzymatic modification of the glycocalyx. Desialylation of adult human cholangiocytes lowers their resistance to toxicity induced by both GCDC and CDC (10, 11). As previously described (10, 11), both GCDC and CDC have high pKas and both exerted toxic effects on adult EHBD cholangiocytes at pH<7.0, which sustained further damage if the glycocalyx had been enzymatically modified. Collectively, TEM and lectin staining highlighting the immature state of the glycocalyx in neonatal mice and functional data on the protective role of the glycocalyx (Supplemental Fig. 4C) offer a compelling mechanism for neonatal susceptibility to injury.

Intrahepatic and extrahepatic cholangiocytes are developmentally distinct (1–3) and the IHBD develops later than the EHBD. Previous reports described immature cell-cell junctions in intrahepatic cholangiocytes in mice as late as 1 week postnatal (29, 30). Our TEM studies show that neonatal IHBD cholangiocytes are compact and well organized and that cell-cell junctions appear similar to those in adults. Neonatal EHBD cholangiocytes, however, have loosely-formed cell-cell junctions and gaps running between them. Our results are based on TEM analysis of intact tissues from BALB/c mice while previous studies were performed using primary cell lines isolated from neonatal and adult C57BL/6 mouse livers (30). Our findings and the published work are consistent, however, in showing higher permeability of cholangiocyte monolayers from neonatal compared to adult cells. As with the glycocalyx, this suggests increased susceptibility to bile acid-mediated damage in the neonatal duct, although in this case primarily the EHBD.

Neonatal EHBD lack collagen and elastin, suggesting that the mechanical properties of the neonatal duct differ from the adult duct. In arteries, elastin and collagen provide resistance to stretch at low and high pressures, respectively, and data from knockout mice suggest that elastin is also necessary for reducing energy loss during loading and unloading (31). Our data thus imply that neonatal ducts have a decreased ability to respond appropriately to mechanical stresses such as that resulting from obstruction.

Small leucine-rich proteoglycans (such as lumican, decorin and fibromodulin) and large proteoglycans (such as versican) regulate collagen fibrillogenesis (32–36). We show here that these proteoglycans as well as HA are present in the EHBD during the postnatal deposition of collagen I. Only lumican and fibromodulin, however, demonstrate altered expression during the postnatal development of the EHBD. Lumican and fibromodulin are also expressed in a sequential manner in developing tendons (33); knock out and in vitro studies suggest that the two proteoglycans mediate the stabilization and destabilization of intermediates necessary for fibril growth into mature bundles (33, 37), raising the possibility that these proteoglycans play a similar role in the development of the EHBD.

The EHBD submucosa is an interstitial space with active fluid flow (12). Here we show that, in an in vitro model of the submucosal matrix, increasing concentrations of collagen in HA lead to a reduction in the diffusion of bile. Although this model has limitations (including that the collagen was non-aligned and the ECM lacked the complexity of the submucosa in vivo), it suggests that bile leakage into the submucosa is less contained and therefore potentially more damaging in the neonate than the adult. Thus, we suggest the immature biliary mucosa (with a sparse glycocalyx and leaky cell-cell junctions) may be mechanistically linked to the immature submucosa (lacking collagen and elastin), the combination leading to bile acid-induced injury, potentially explaining in part neonatal *susceptibility* to injury.

The neonatal submucosa may also be important in the *response* to bile duct injury. The cells of the EHBD submucosa and their responses to bile acids or other agents have not been well characterized. We previously showed that in the human adult, these cells are CD34, vimentin and D2-40 positive but do not stain for lymphovascular (CD31, ERG or LYVE-1) or stem cell (CD117 or nuclear β catenin) markers, suggesting that they are either fibroblasts or a form of mesenchymal stem cells (12). Here, we have confirmed that in the mouse, cells in the submucosal space are mostly fibroblasts (staining positive for CD34, vimentin and PDGFRβ). The population appears heterogeneous, however: there are also αSMA-negative, CD105-negative and vimentin/CD34-positive cells in the adult. Larger percentages of α-SMA-negative and CD105-positive cells are present in neonatal than in adult EHBDs. Additionally, in contrast to cells in the adult submucosa, the majority of cells in the neonatal submucosa cells are actively producing collagen. This suggests that neonatal submucosal cells have high fibrogenic potential and raises the interesting possibility that injury to the neonatal EHBD occurs in an environment that is primed for fibrosis. If these cells persist in a quiescent state in the adult, it will be important to determine whether they reactivate after injury, in particular in sclerotic biliary diseases such as primary sclerosing cholangitis with its classical “onion-skin fibrosis” or cholangiocarcinoma.

Taken as a whole, our data show that neonatal bile ducts, especially the EHBD, are anatomically and functionally immature – with a sparse glycocalyx, leaky cholangiocyte cell-cell junctions, a structurally undeveloped submucosal matrix, and actively collagen-secreting cells – such that they are likely to be highly susceptible to injury and fibrosis. This adds to our understanding of potential mechanisms of BA and other forms of neonatal bile duct injury (11), and may have implications for understanding duct injury in adult sclerosing biliary diseases as well.

## Authors’ contributions

G.K & J.L. contributed to study concept and design, acquisition, analysis, and interpretation of data, and drafting of the manuscript. A.K., Y.D., R.G. & X.L. contributed to acquisition, analysis, and interpretation of data. O.W.-Z. & N.J contributed to acquisition of data and study concept and design. T.K. provided material support and contributed to data interpretation. P.A.R. provided material support and contributed to analysis and interpretation of data. N.D.T. contributed to study concept and design, and provided data interpretation and critical revision of the manuscript. R.G.W. conceived ideas and designed the research, provided critical revision of the manuscript, obtained funding and supervised the study.

## Abbreviations

IHBD: intrahepatic bile duct
EHBD: extrahepatic bile duct
BA: biliary atresia
DAB: diaminobenzidine
PBS: phosphate buffered saline
SMA: smooth muscle actin
DAPI: 4’,6-diamidino-2-phenylindole
TEM: transmission electron microscopy
SHG: second harmonic generation
Con A: Concanavalin A
PHA-E: Phaseolus vulgaris erythroagglutinin
PHA-L: Phaseolus vulgaris Leucoagglutinin
PNA: Peanut Agglutinin
GSL: Griffonia (Bandeiraea) simplicifolia
SBA: Soybean
MAI II: Maachia ammurensis II
HA: hyaluronic acid
RER: rough endoplasmic reticulum
GCDC: glycochenodeoxycholic acid

## Acknowledgments

We thank Richard Gomer (Texas A&M) for advice on lectin staining, Robert Heuckeroth (UPenn) for advice on nerve staining, and Karin Diggle (UCSD) for assistance with animal work. We gratefully acknowledge the assistance of the following core facilities and individuals at the University of Pennsylvania: the Penn Vet Imaging Core and Gordon Ruthel, the Perelman School of Medicine Cell and Developmental Biology Microscopy Core and Electron Microscopy Resource Laboratory and Andrea Stout, and the NIDDK Center for Molecular Studies in Digestive and Liver Diseases Molecular Pathology and Imaging Core (NIH P30 DK050306).

## Supplemental Methods

### Histology, immunostaining, and lectin staining

For paraffin-embedded tissue, samples were dewaxed with xylene, then rehydrated through a graded series of alcohols and distilled water before undergoing heat-mediated antigen retrieval in 10 mM citric acid buffer (pH 6.0). Frozen sections were brought to room temperature before fixation in 10% neutral buffered formalin for 4 min. For diaminobenzidine (DAB) labelling, sections were incubated in 3% H_2_O_2_ to quench endogenous peroxidases, then blocked with StartingBlock™ T20/phosphate buffered saline (PBS) Blocking Buffer (Thermo Fisher Scientific, Waltham, MA) or blocking buffer containing 10% serum and 1% BSA and then were incubated in primary antibodies (0.2% Triton X-100, 0.1% bovine serum albumin, in PBS) overnight at 4°C. Antibodies used are listed in Supplemental Tables S1 and S2. Hyaluronic acid was stained on frozen sections using biotinylated hyaluronic acid binding protein (Millipore, Burlington, MA; 385911). Cy2-, Cy3-and Cy5-conjugated secondary antibodies were used (1:500, Vector Laboratories, Burlington, CA).

For lectin staining, endogenous biotin was blocked by incubating in streptavidin and biotin solutions for 15 min followed by blocking in Carbo-Free Blocking Solution for 1 h. The slides were then incubated with biotinylated lectins for 1 h at room temperature followed by brief washes in 1XPBS. Biotin was then detected by Streptavidin, Alexa Fluor™ 488 conjugate (Invitrogen, Carlsbad, CA). Sections were mounted in medium containing 4’,6-diamidino-2-phenylindole (DAPI; Vector Laboratories). Stained sections were imaged using a SCTR Leica confocal microscope and Leica application suite (LAS X) (Leica, Buffalo Grove, IL) and the raw image files were processed using Fiji ImageJ software.

For Alcian Blue staining, slides were deparaffinized and hydrated as above, then soaked in Acetic Acid 3% aqueous solution (Poly Scientific R&D Corp., Bay Shore, NY) for 3 min followed by 1% Alcian Blue in 3% Acetic Acid (Poly Scientific R&D Corp.), pH 2.5 for 30 min. Slides were rinsed in dH_2_O and counterstained with Nuclear Fast Red 0.1% (Poly Scientific R&D Corp.) for 40 sec, then rinsed in dH_2_O, dehydrated in ethanol and xylene and mounted in Cytoseal 60 (Thermo Fisher Scientific). Samples were imaged using a Nikon Eclipse E600 microscope attached to a Digital Sight DS-U3 microscope camera controller (Nikon Instruments Inc., Melville, NY). Images were processed using NIS-Elements BR (Nikon Instruments Inc.) imaging software.

**Supplemental Table 1:** Antibodies used on paraffin embedded tissue

| Antigen | Concentration/<br>Dilution | Source |
| --- | --- | --- |
| Acetyl-alpha Tubulin<br>(Lys40) | 1-3 µg/ml | Thermofischer<br>Scientific, 32-2700 |
| αSMA | 1:100 | Sigma, A2547 |
| Caspase 3 | 1:500 | Sigma, C8487 |
| CD34 | 1:100 | Abcam, ab81289 |
| K19 | 1:100 | DSHB, Troma III |
| Collagen I | 1:50 | Southern Biotech,<br>1310-01 |
| Collagen III | 1:300 | Abcam, ab7778 |
| Elastin | 1:200 | BIOSS, 1756R |
| Ki67 | 1:100 | Abcam, ab16667 |
| Lumican | 1:200 | Abcam, ab168348 |
| Lyve-1 | 1:100 | Abcam, ab14917 |
| PDGFR β | 1:100 | Cell signaling, 3169 |
| TuJ1 (β-Tubulin III) | 1:10 000 | Covance, PRB-435P |
| Vimentin | 1:500 | Novus, NB300 223 |

**Supplemental Table 2:** Antibodies used on frozen tissue

| Antigen | Dilution | Source |
| --- | --- | --- |
| Aggrecan | 1:200 | Thermo scientific,<br>MA3-16888 |
| αSMA | 1:100 | Sigma, A2547 |
| CD105 | 1:50 | R&D, MAB1320 |
| CD34 | 1:100 | Abcam, ab81289 |
| Collagen 4 | 1:100 | Abcam, ab6586 |
| Decorin | 1:50 | Abcam, ab175404 |
| F4/80 | 1:50 | Abcam, ab6640 |
| Fibromodulin | 1:200 | Abcam, ab81443 |
| Fibronectin | 1:50 | Abcam, ab2413 |
| GFP | 1:1000 | Abcam, ab13970 |
| Versican | 1:50 | EMD Millipore,<br>AB1033 |
| Vimentin | 1:500 | Novus, NB300 223 |

**Supplemental Table 3:** Lectin panel

| **Lectin** | **Concentration** | **Source** |
| --- | --- | --- |
| *Phaseolus Vulgaris*<br>*Leucoagglutinin* (PHA-L) | 1 µg/ml | Vector<br>Laboratories Inc,<br>B-1115 |
| *Phaseolus Vulgaris*<br>*Erythroagglutinin* (PHA-E) | 1 µg/ml | Vector<br>Laboratories Inc,<br>B-1125 |
| *Concanavalin A* (ConA) | 1 µg/ml | Vector<br>Laboratories Inc,<br>B-1005 |
| *Griffonia (Bandeiraea)*<br>*simplicifolia lectin* (GSL) | 1 µg/ml | Vector<br>Laboratories Inc,<br>B-1205 |
| *Glycine max* (soybean)<br>*agglutinin* (SBA) | 1 µg/ml | Vector<br>Laboratories Inc,<br>B-1015 |
| *Arachis hypogaea* (peanut)<br>*agglutinin* (PNA) | 1 µg/ml | Vector<br>Laboratories Inc,<br>B-1075 |
| *Maackia Amurensis Lectin II*<br>(MALII)[WR16] | 1 µg/ml | Vector<br>Laboratories Inc,<br>B-1265 |
| *Lens culinaris agglutinin* (LCA) | 1 µg/ml | Vector<br>Laboratories Inc,<br>B-1045 |
| *Pisum sativum agglutinin* (PSA) | 1 µg/ml | Vector<br>Laboratories Inc,<br>B-1055 |
| *Sambucus Nigra Lectin* (SNA) | 1 µg/ml | Vector<br>Laboratories Inc,<br>B-1305 |
| *Succinylated Wheat Germ*<br>*Agglutinin* (WGA) | 1 µg/ml | Vector<br>Laboratories Inc,<br>B-1025S |
| *Sophora japonica* (SJA) | 1 µg/ml | Gift from Gomer<br>lab |

**Supplemental Fig. 1.**
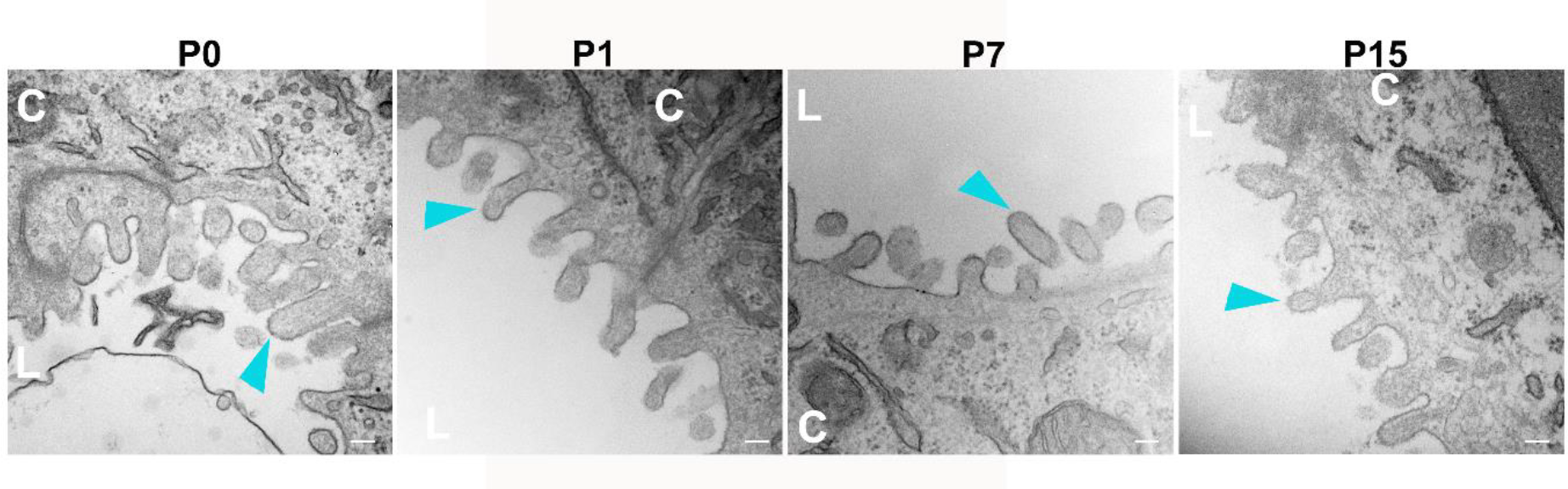
The neonatal IHBD lacks a mature glycocalyx. TEM images of BALB/c mouse livers at postnatal days 0 (P0), 1 (P1), 7 (P7) and 15 (P15). Magnification ×75,000. Representative images shown of livers from 3 animals stained per each age group. Teal arrowheads point to the glycocalyx. Scale bars, 500 nm. **C** denotes cholangiocytes and **L** lumen.

**Supplemental Fig. 2.**
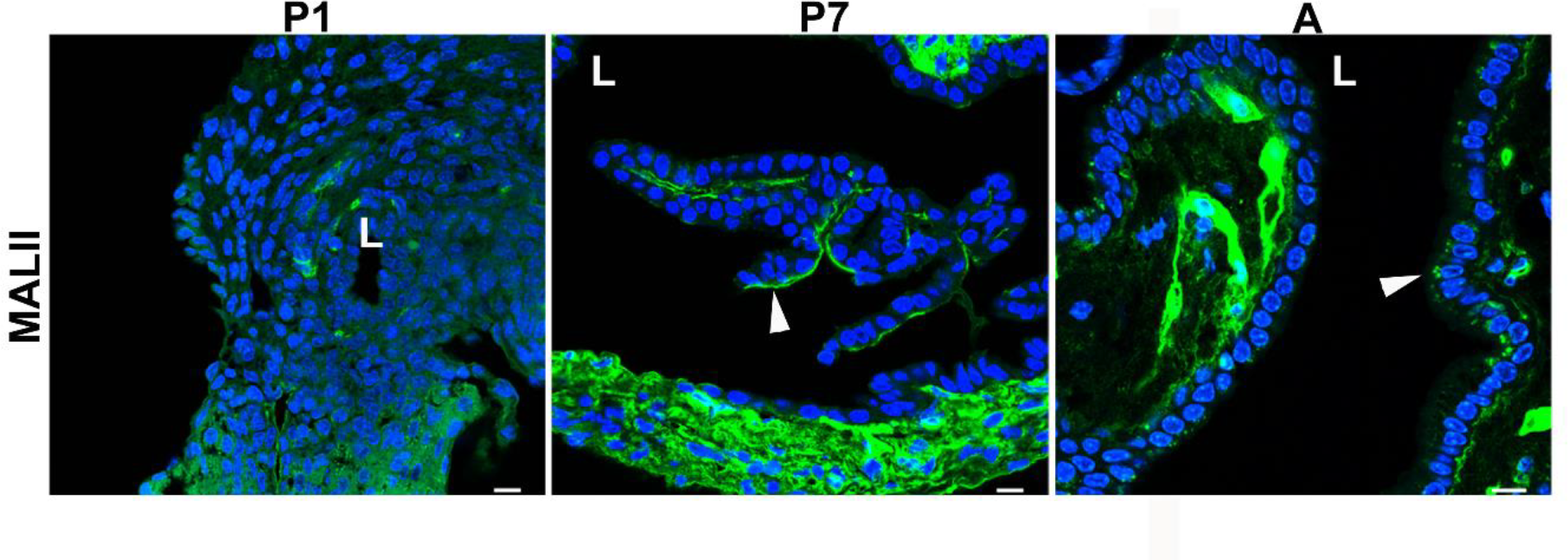
MALII lectin staining of BALC/c mouse EHBDs varies during EHBD development. EHBDs from mice at postnatal day 1 (P1) and day 7 (P7) and from adults (A) were stained with MALLII lectin (green) and DAPI (nuclei; blue). White arrowheads point to the stained glycocalyx. Representative images shown from 3 stained EHBD per age. Scale bars, 10 μm. **L** denotes lumen.

**Supplemental Fig. 3.**
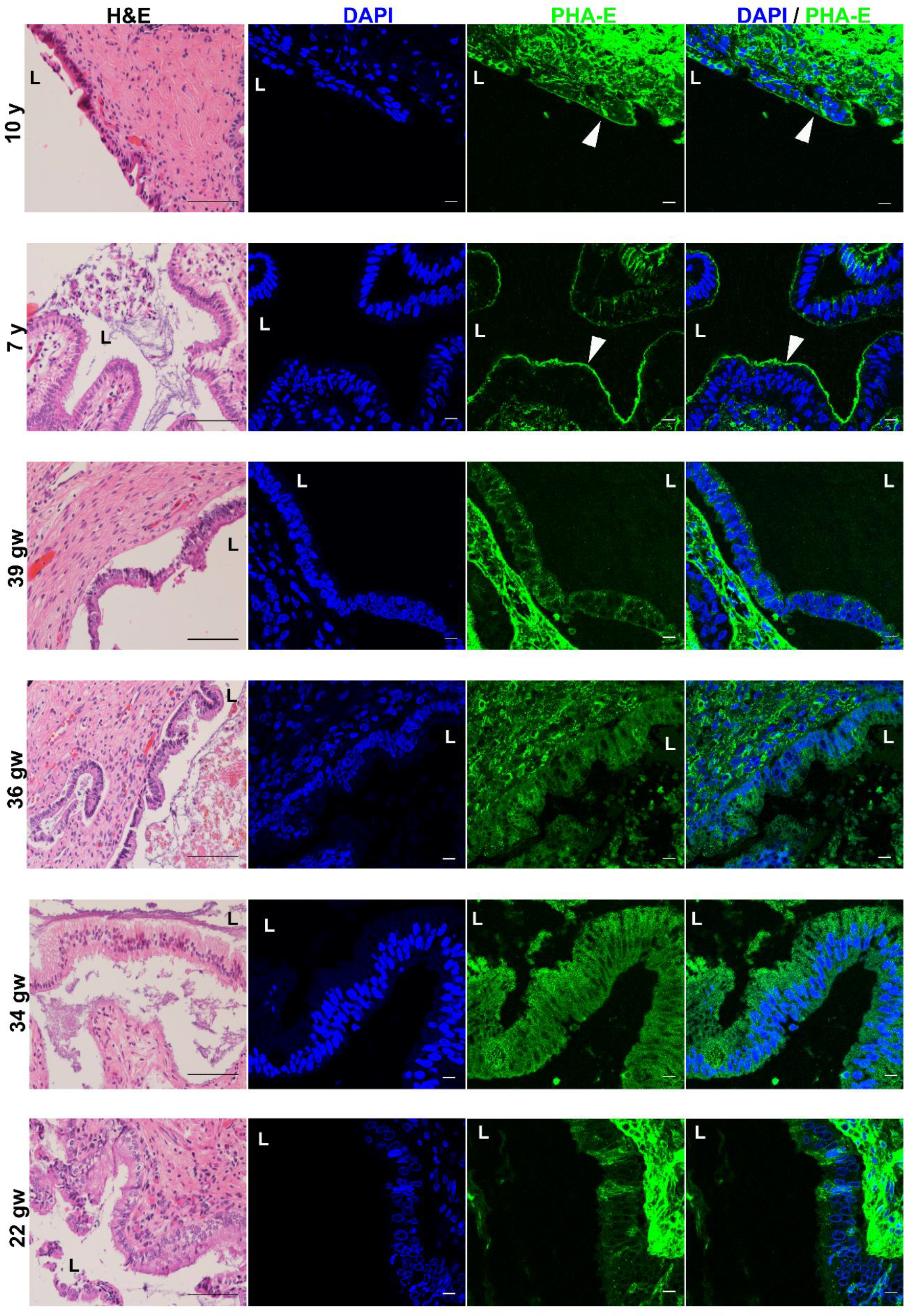
PHA-E lectin staining of human EHBD sections at 22, 34, 36, 39 gestational weeks (gw) and 7 and 10 years (y) of age. PHA-E lectin (green), nuclei stained with DAPI (blue). White arrowheads point to the stained glycocalyx, which is not seen in the prenatal specimens. Representative images from each sample shown. Scale bars, 10 μm. Corresponding H&E staining shown on left. Scale bar, 100 µm. **L** denotes lumen.

**Supplemental Fig. 4.**
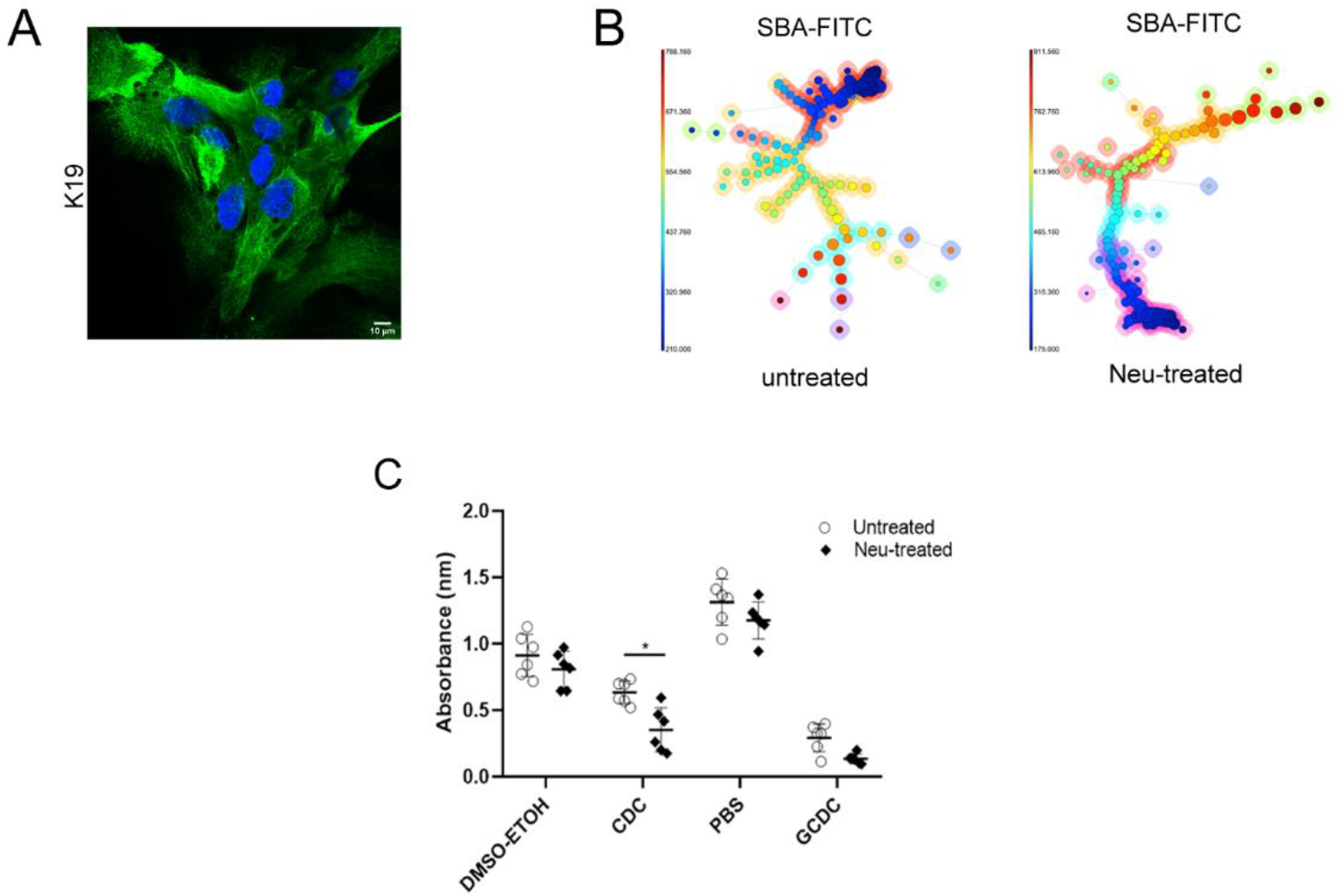
Neonatal cholangiocyte spheroids are susceptible to bile acid-induced injury. A) Immunostaining of adult primary cholangiocytes using K19 antibody (green). B) Minimal spanning tree visualization of adult EHBD cholangiocytes by FlowSOM analysis. Manual gating results for FITC expression. An increase in FITC labelling was observed after neuraminidase treatment, indicating successful removal of sialic acid residues. C) Metabolic activity measurements using the WST-1 assay after exposure of neuraminidase (neu)-treated and control primary cholangiocytes isolated from adult BALB/c mice to the bile salts CDC and GCDC. All results are expressed as means ± SD of at least 3 independent experiments. Data shown are mean ± SD, * P<0.05.

**Supplemental Fig. 5.**
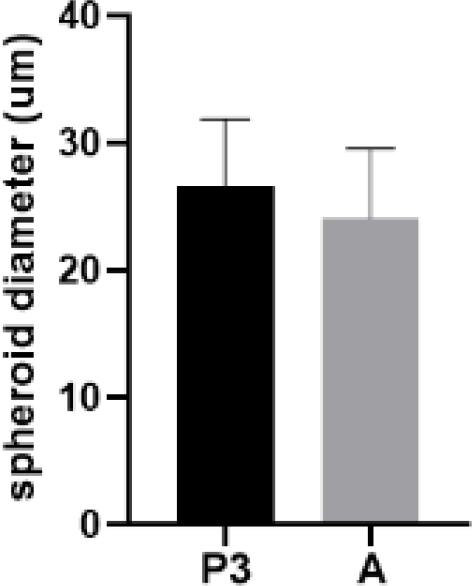
Spheroid size in neonatal and adult cholangiocytes. The mean diameter of the spheroids made from P3 cholangiocytes (N=25) was 26 μm while that of adult cholangiocytes (N=26) was 24 μm. Data shown are mean ± SD (t-test, ** P=0.1044, no significant difference).

**Supplemental Fig. 6.**
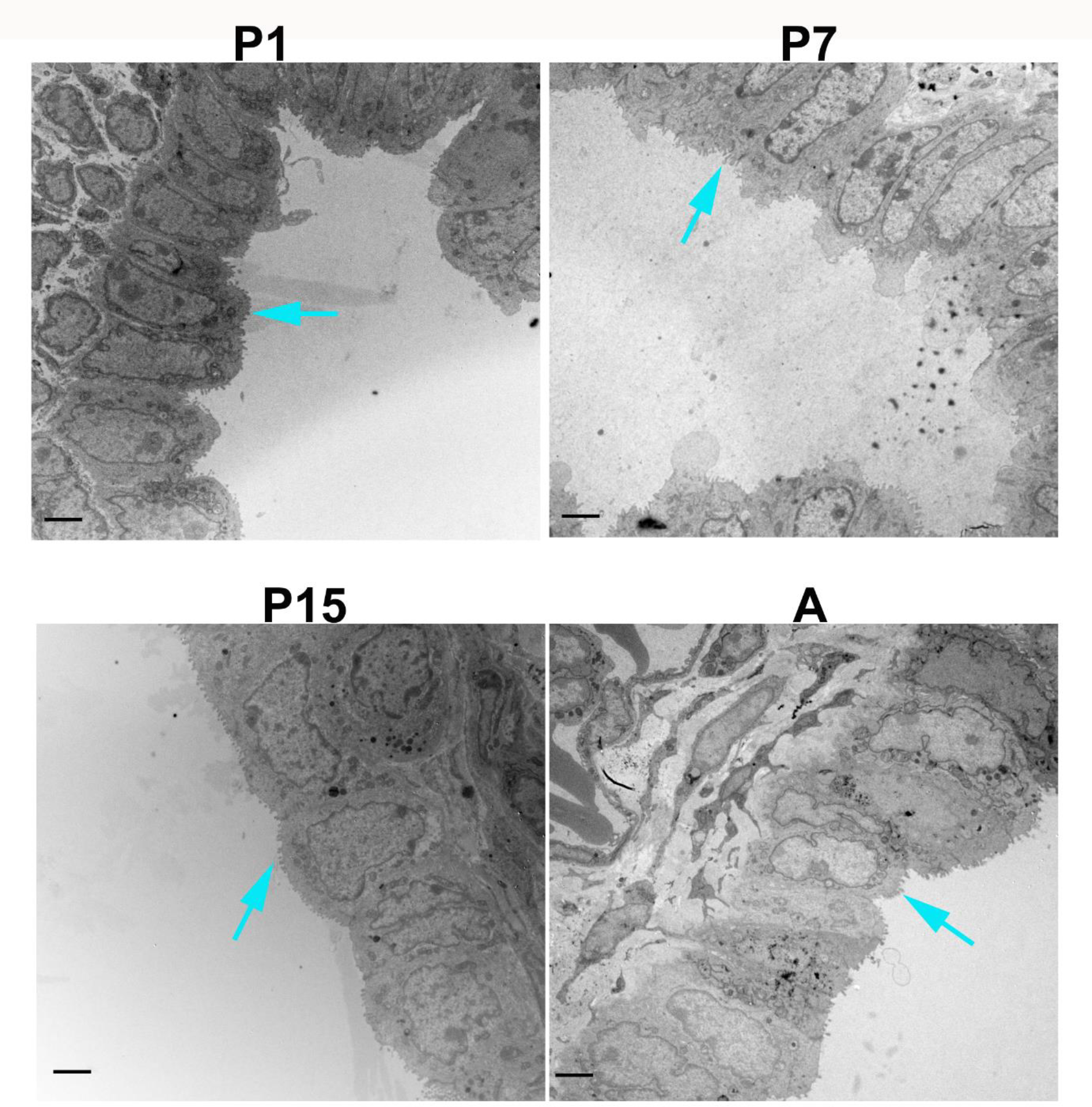
Microvilli are abundant on the luminal side of BALB/c cholangiocytes. TEM images of cholangiocytes from BALB/c EHBDs at postnatal days 1 (P1), 7 (P7), 15 (P15) and adult (A). Magnification X 5000. Teal arrows point to microvilli. Representative images shown of EHBDs from 3 animals per age group. Scale bars, 2 μm.

**Supplemental Fig. 7.**
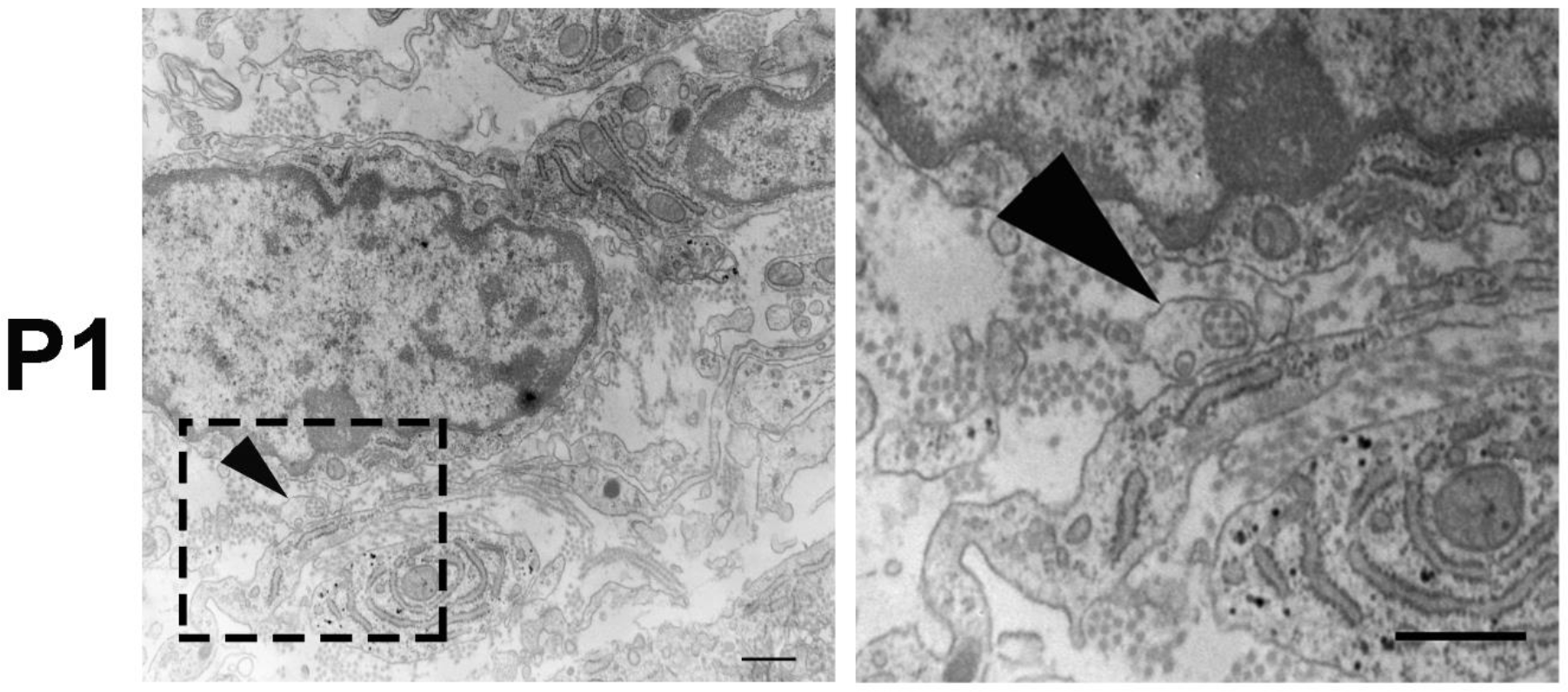
Fibripositor-like structure found in neonatal EHBDs. TEM image of an EHBD from a P1 pup. Arrow points to fibripositor-like structure. Scale bar, 500 nm.

**Supplemental Fig. 8.**
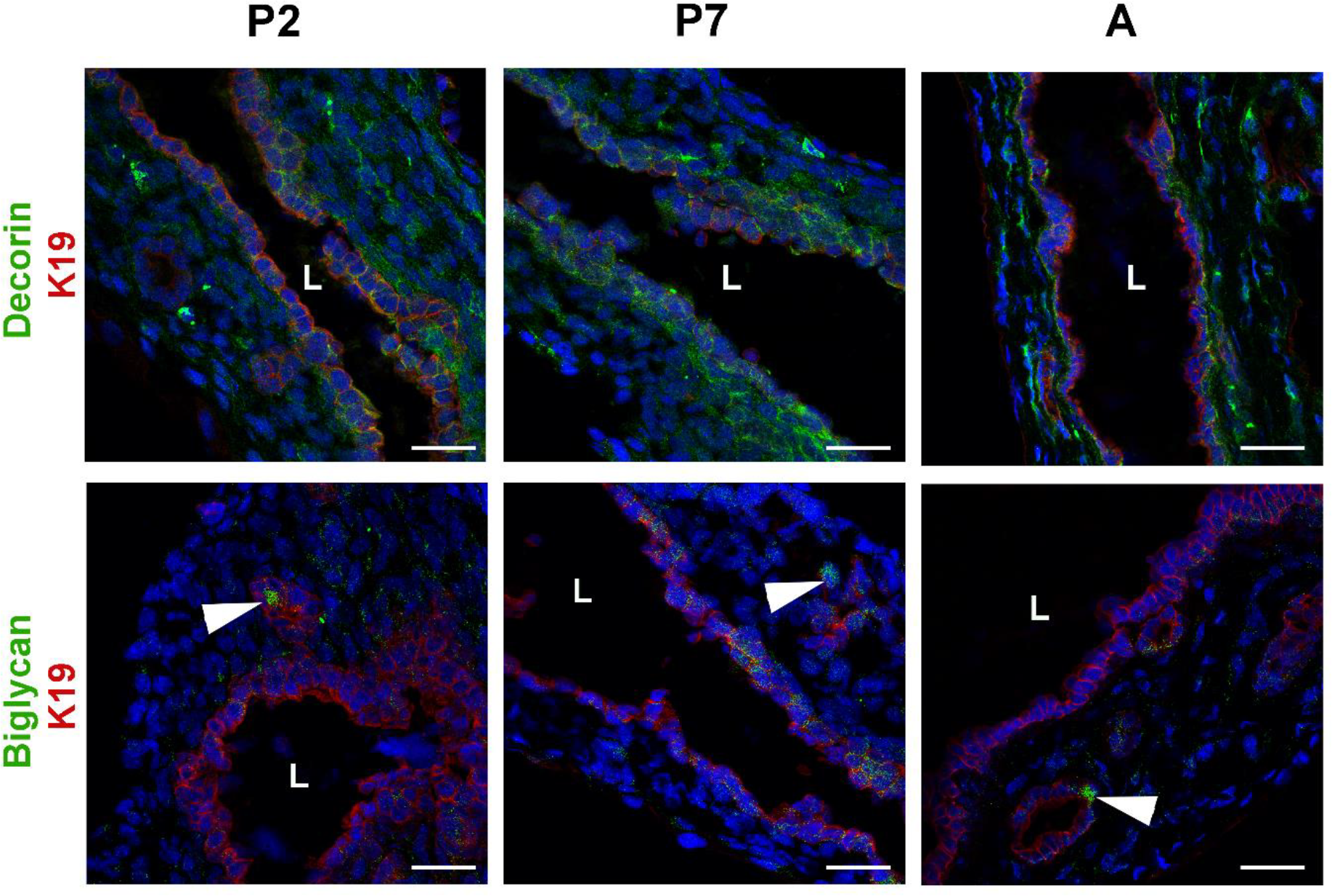
Decorin and biglycan are present at low levels from birth. EHBD from 2 (P2) and 7 (P7) day old pups as well as adult (A) mice were stained for proteoglycans decorin (green) and biglycan (green). Nuclei were stained with DAPI (blue). Representative images shown of EHBDs from 3-4 animals stained per age group. Scale bars, 100μm. **L** denotes lumen.

**Supplemental Fig. 9.**
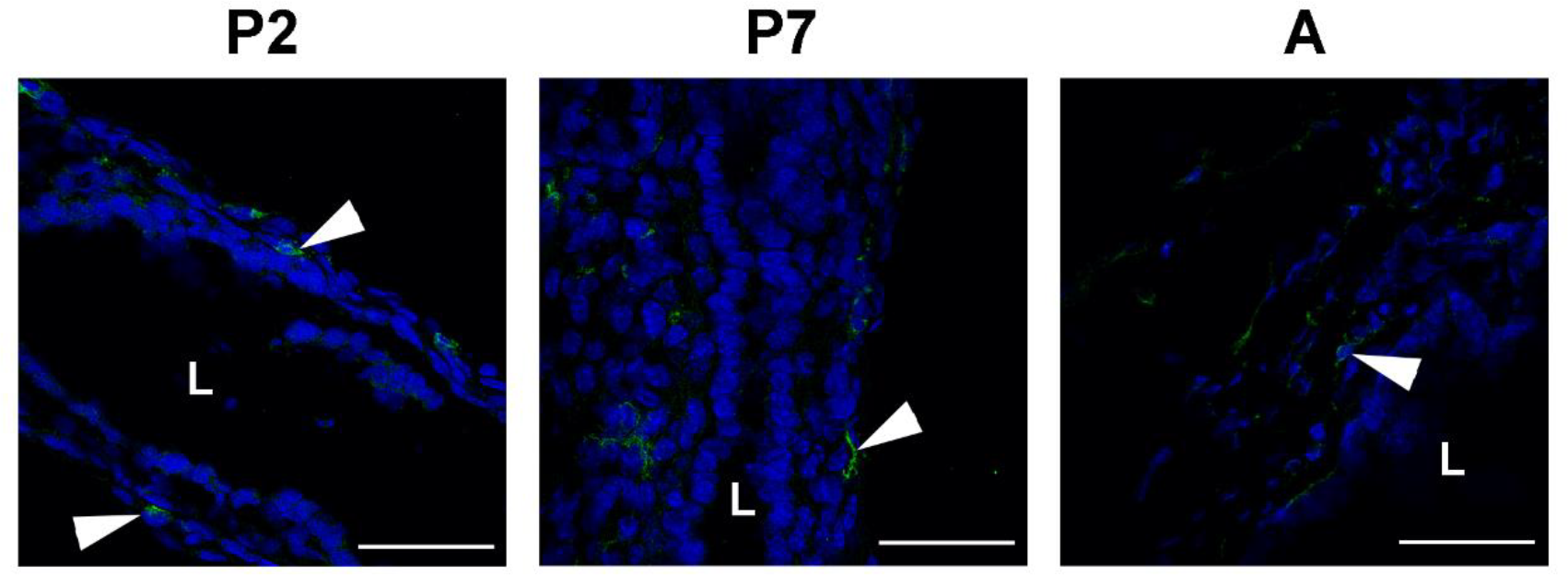
Macrophages are found within the mouse EHBD submucosa. EHBD from 2 (P2) and 7 (P7) day old pups as well as adult (A) mice were stained with the macrophage marker F4/80 (green), indicated by white arrowheads. Nuclei were stained with DAPI (blue). Representative images shown of EHBDs from 3-4 animals stained per age group. Scale bars, 100 μm. **L** denotes lumen.

**Supplemental Fig. 10.**
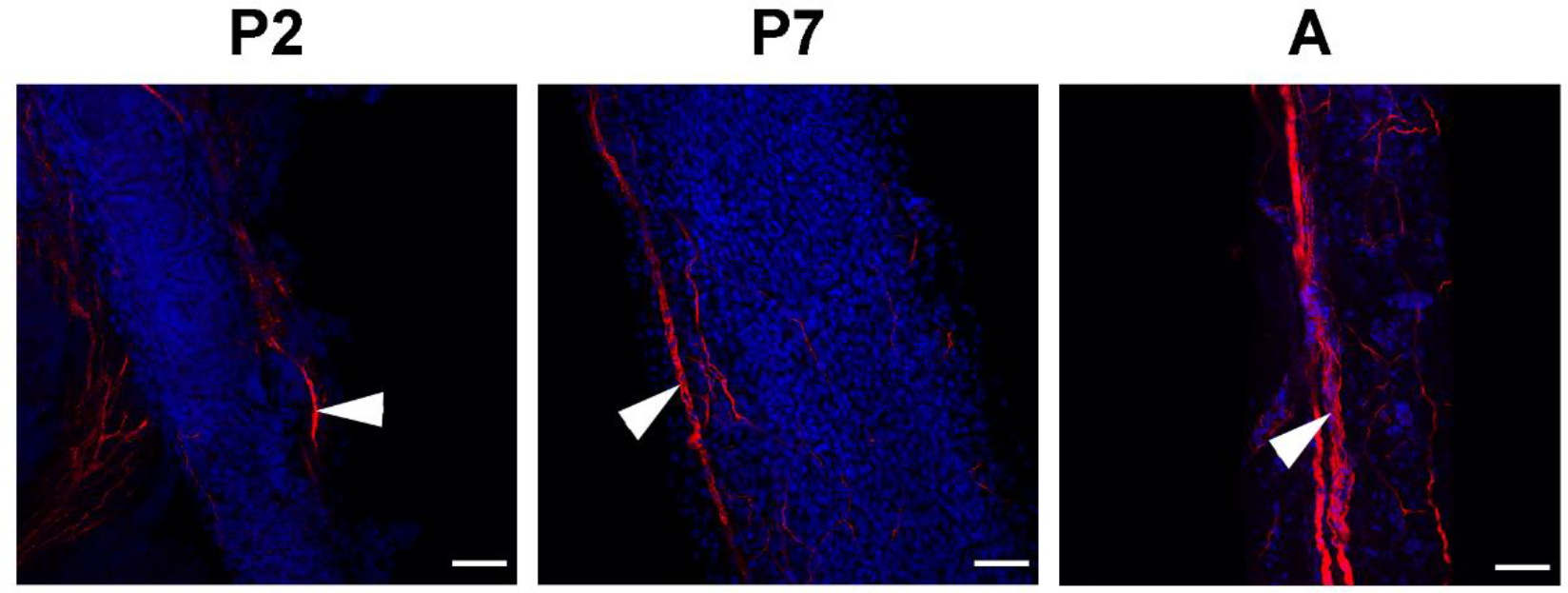
Mouse EHBDs are innervated from birth. The submucosa of EHBDs from postnatal day 2 (P2) and 7 (P7) mice as well as adults (A) were stained with the neuronal marker neuron-specific class III β-tubulin (TUJI, red), indicated by the white arrowheads. Nuclei were stained with DAPI (blue). Representative images of stains carried out on 3-4 EHBD per age group are shown. Scale bars, 100 μm.

## Notes

**Conflict of interest**: The authors declare no conflicts of interest that pertain to this work.

**Grant Support:** Supported in part by National Institutes of Health grants DK092111 and DK119290 (to R.G.W.), the Fred and Suzanne Biesecker Pediatric Liver Center at The Children’s Hospital of Philadelphia. This work was also supported by the Center for Engineering MechanoBiology (CEMB), an NSF Science and Technology Center, under grant agreement CMMI:15-48571. A.K. was a recipient of a fellowship through the CHOP Training Program in the Genetic Basis of Pediatric Gastrointestinal Disorders (5T32DK101371). O.W.Z. was a recipient of a fellowship grant through the American Association for the Study of Liver Diseases (AASLD).

